# In defense of the original Type I functional response: The frequency and population-dynamic effects of feeding on multiple prey at a time

**DOI:** 10.1101/2024.05.14.594210

**Authors:** Mark Novak, Kyle E. Coblentz, John P. DeLong

**Author notes:** **Code and data availability** The FoRAGE compilation is available from the *Knowledge Network for Biocomplexity* (DeLong & Uiterwaal, 2018). All code and data are available at https://github.com/marknovak/FR_n-prey-at-a-time and DOI: 10.6084/m9.figshare.28292147 (Novak et al., 2025a;b). **Author contributions** MN conceived of the study, performed the analyses, and wrote the first draft. JPD compiled functional response datasets. KEC and JPD discussed the analyses and edited the manuscript.

## Abstract

Ecologists differ in the degree to which they consider the linear Type I functional response to be an unrealistic versus sufficient representation of predator feeding rates. Empiricists tend to consider it unsuitably non-mechanistic and theoreticians tend to consider it necessarily simple. Holling’s original rectilinear Type I response is dismissed by satisfying neither desire, with most compromising on the smoothly saturating Type II response for which searching and handling are assumed to be mutually exclusive activities. We derive a “multiple-prey-at-a-time” response and a generalization that includes the Type III to reflect predators that can continue to search when handling an arbitrary number of already-captured prey. The multi-prey model clarifies the empirical relevance of the linear and rectilinear models and the conditions under which linearity can be a mechanistically-reasoned description of predator feeding rates, even when handling times are long. We find support for linearity in 35% of 2,591 compiled empirical datasets and support for the hypothesis that larger predator-prey body-mass ratios permit predators to search while handling greater numbers of prey. Incorporating the multi-prey response into the Rosenzweig-MacArthur population-dynamics model reveals that a non-exclusivity of searching and handling can lead to coexistence states and dynamics that are not anticipated by theory built on the Type I, II, or III response models. In particular, it can lead to bistable fixed-point and limit-cycle dynamics with long-term crawl-by transients between them under conditions where abundance ratios reflect top-heavy food webs and the functional response is linear. We conclude that functional response linearity should not be considered empirically unrealistic but also that more cautious inferences should be drawn in theory presuming the linear Type I to be appropriate.

## Introduction

The way that predator feeding rates respond to changes in prey abundance, their functional response, is key to determining how species affect each other’s populations (Murdoch & Oaten, 1975). The challenge of empirically understanding and appropriately modeling functional responses is therefore central to myriad lines of ecological research that extend even to the projection of Earth’s rapidly changing climate (DeLong, 2021; Rohr *et al*., 2023).

The simplest functional response model, the Type I response, describes feeding rates as increasing linearly with prey abundance. Interpreted to represent an analytically-tractable first-order approximation to all other prey-dependent forms (Lotka, 1925; Volterra, 1926), its simplicity has caused the Type I to become foundational to theory across Ecology’s many subdisciplines. Nonetheless, there is a common and persistent belief among empirically-minded ecologists that the Type I response is unrealistic and artifactual. Indeed, it is typically dismissed *a priori* from both empirical and theoretical efforts to “mechanistically” characterize predator feeding rates (e.g., Baudrot *et al*., 2016; Kalinkat *et al*., 2023). This dismissal is similarly levied at the piecewise rectilinear response (e.g., Koen-Alonso, 2007), originally referred to by Holling (1959a) as the Type I response (Denny, 2014; Holling, 1965), in which feeding rates increase linearly with prey abundance to a relatively abrupt maximum. Support comes from syntheses concluding functional response linearity to be rare, with feeding rates more consistent with smoothly saturating Type II responses being by far the more frequently inferred (Dunn & Hovel, 2020; Jeschke *et al*., 2004).

Countering justifications for the continued use of the linear Type I response in theory relate to the challenge of extrapolating the inferences of mostly small-scale experiments to natural field conditions (DeLong, 2021; Griffen, 2021; Jeschke et al., 2004; Li et al., 2018; Novak & Stouffer, 2021b; Novak *et al*., 2017; Uiterwaal et al., 2018). For example, prey abundances in the field may vary relatively little over relevant scales, making linearity a sufficiently good approximation for how species affect each other (Wootton & Emmerson, 2005). Further, prey abundances in nature are typically much lower than those used in experiments to elicit predator saturation (Coblentz *et al*., 2023), which may consequently be rare in nature (but see Jeschke, 2007). Functional responses could therefore be approximately linear even for predator-prey interactions having very long handling times (e.g., Novak, 2010).

Here, our goal is to offer a further way of resolving ecologists’ views on the linear and rectilinear models by considering a reason for feeding rates to exhibit linear prey dependence over a large range of prey abundances. This reason is not one of experimental design or variation in prey abundances per se, but rather is attributable to the mechanics of predator-prey biology: the ability of predator individuals to handle and search for more than just one prey individual at a time (i.e. the non-exclusivity of handling and searching). Although it is straightforward to show how the linear Type I can emerge when handling times are assumed to be entirely inconsequential, and although functional response forms that could result from a non-exclusivity of handling and searching have been considered before (Jeschke *et al*., 2002; 2004; Mills, 1982; Sjöberg, 1980; Stouffer & Novak, 2021), we contend that the empirical relevance and potential prevalence of such “multiple-prey-at-a-time” feeding (henceforth multi-prey feeding) are not sufficiently understood due to an inappropriately literal interpretation of the “handling time” parameter of functional response models (see *Discussion* and DeLong, 2021; Jeschke et al., 2002; 2004). Likewise, the potential implications of multi-prey feeding for predator-prey coexistence and population dynamics have not, to our knowledge, been assessed.

We begin by providing a derivation of a simple multi-prey functional response model for a single predator population feeding on a single prey species that relaxes the assumption of searching and handling being exclusive activities. This derivation helps clarify the empirical relevance of the linear and rectilinear models and the conditions under which these can be good descriptions of feeding rates (Jeschke *et al*., 2004). We then further generalize the multi-prey model to include the Holling-Real Type III response and fit all models to a large number of datasets assembled in a new version of the FoRAGE compilation (Uiterwaal *et al*., 2022). This allows us to quantify the potential prevalence of multi-prey feeding and to test the hypothesis that larger predator-prey body-mass ratios permit predators to handle and search for more prey at a time. We also assess the predicted association between larger body-mass ratios and more pronounced Type III responses. Finally, we incorporate the multi-prey response into the Rosenzweig & MacArthur (1963) “paradox of enrichment” population-dynamic model to assess its potential influence on predator-prey coexistence and dynamics.

With our statistical analyses demonstrating that many datasets are indeed consistent with multi-prey feeding and that larger predator-prey body-mass ratios are indeed more conducive to multi-prey feeding (and more pronounced Type III responses), our mathematical analyses demonstrate that even small increases in the number of prey that a predator can handle at a time can lead to dynamics that are not anticipated by theory assuming Type I, II, or III response models.

### A functional response for multi-prey feeding

#### Holling’s Type II response

The multi-prey model may be understood most easily by a contrast to Holling’s Type II model (a.k.a. the disc equation, Holling, 1959b). There are several ways to derive the Type II (Garay, 2019), but the most common approach takes the perspective of a single predator individual that can either be searching or “handling” a single prey individual at any point in time: In the time *T*_*S*_ that a predator spends searching it will encounter prey at a rate proportional to their abundance *N*, thus the number of prey eaten is *N*_*e*_ = *aNT*_*S*_ where *a* is the attack rate. Rearranging we have *T*_*S*_ = *N*_*e*_*/aN* . With a handling time *h* for each prey, the length of time spent handling all eaten prey will be *T*_*H*_ = *hN*_*e*_. Given the presumed mutual exclusivity of the two activities, *T*_*S*_ = *T - T*_*H*_ where *T* is the total time available. Substituting the second and third equations into the fourth, it follows that *N*_*e*_ = *aNT/*(1 + *ahN*). We arrive at the predator individual’s feeding rate by dividing by *T*, presuming steady-state predator behavior and constant prey abundances.

An alternative derivation on which we build to derive the multi-prey model considers a temporal snapshot of a predator population composed of many identical and independent individuals (see also Real (1977) and the *Supplementary Materials*). Assuming constant prey abundance and steady-state conditions, the rate at which searching individuals *P*_*S*_ become handling individuals *P*_*H*_ must equal the rate at which handling individuals become searching individuals such that 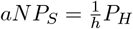,visually represented as

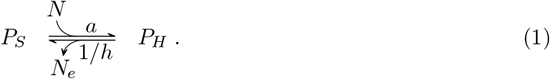

Given the mutual exclusivity of searching and handling, *P*_*S*_ = *P - P*_*H*_, where *P* is the total number of predators. Substituting this second equation into the first, it follows that the total number of handling predators *P*_*H*_ = *ahNP/*(1 + *ahN*). Eaten prey are generated at rate 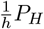 by all these predators as they revert back to searching. We thus obtain Holling’s Type II (per-predator) model by multiplying the proportion of handling predators, 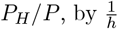.

### The multi-prey response

The derivation of the multi-prey response follows the same logic but assumes that searching and handling are not mutually exclusive activities until an arbitrary count of *n* prey individuals are being handled (see the *Supplementary Materials* for a more explicit derivation); handling need not reflect literal handling but rather could also reflect a process of digestion and stomach fullness.

With constant prey abundance and steady-state conditions as before, we assume that predators continue to handle each prey with handling time *h* and that predators handling less than *n* prey continue to search for and encounter prey at rate *aN* . The rate at which searching individuals *P*_*S*_ become 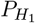 individuals handling one prey is then equal to the rate at which they revert back to being searching individuals with no prey, thus 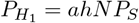.Likewise, the rate at which 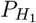 individuals become 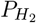 individuals handling two prey must equal the rate these revert back to handling just one prey, thus 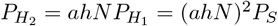.That is,

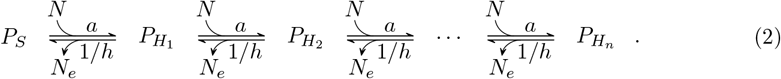

Generalizing by induction, the number of predators *P*_*H*_ handling *i* prey will be (*ahN*)^*i*^*P*_*S*_ for *i* 2 {1, 2, 3,. .., *n*}. The proportion of predators handling *i* prey at any point in time will then be

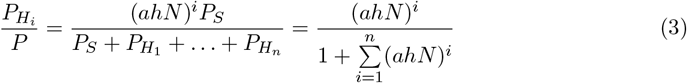

(Fig. S.1). With each of these groups generating eaten prey at rate 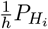,the per predator feeding rate of the population is obtained by a summation across all groups, giving

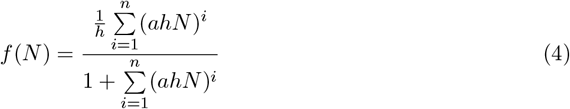

(Fig. 1). This is the multi-prey model for integer values of *n*. However, because the geometric series 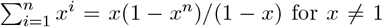 for *x* ≠ 1, we can also write the model more generally for arbitrary values of *n* as

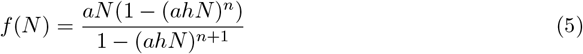

to reflect predator populations capable of searching while handling a non-integer (e.g., average) number of prey individuals.

**Figure 1:**
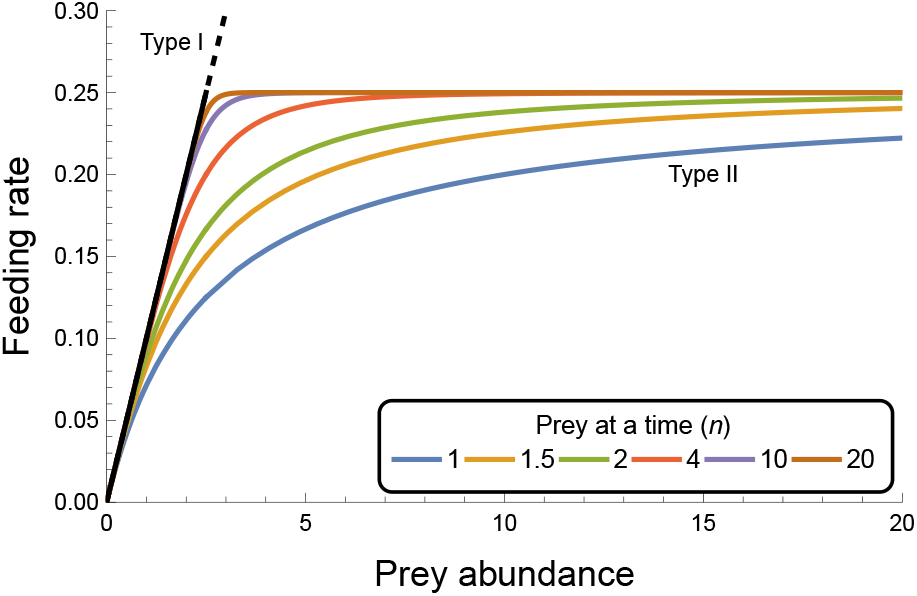
The potential forms of the multi-prey response. The multi-prey model diverges from the Type II (for which *n* = 1) and approaches the rectilinear model as the number *n* of prey individuals that a predator can handle while continuing to search increases. When *n* = ∞ it reduces to the linear Type I which can remain a biologically appropriate description of predator feeding rates so long as *ahN <* 1 (indicated by non-dashed region of the black line). *Parameter values*: attack rate *a* = 0.1 and handling time *h* = 4.

We note that Sjöberg (1980) derived equivalent formulations in Michaelis-Menten enzyme-kinematics form with parameters having correspondingly different statistical properties (Novak & Stouffer, 2021a; Rohr *et al*., 2022). We also note that despite the appearance of two summations in eqn. 4 and the unusual appearance of subtractions in eqn. 5 (see *Supplementary Materials*), the model has only three parameters and thus has a parametric complexity no greater than that of the Holling-Real Type III model and many others (see Table 1 of Novak & Stouffer, 2021a). In fact, for subsequent model-fitting, we will combine the multi-prey and Holling-Real models to a four-parameter generalization,

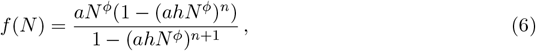

which can be simplified to the other models when *ϕ* = 1. Parameter *ϕ* (a.k.a. the Hill exponent) can be interpreted as the number of prey encounters a predator must experience before its feeding efficiency is maximized (Real, 1977).

### Relevance of the Type I response

The conditions under which the linear, rectilinear, and Type II models can be good descriptions of predator feeding rates are clarified by observing that the multi-prey response simplifies to the Type II when *n* = 1 and approaches the rectilinear model as *n* increases (Fig. 1). Further, the linear Type I is obtained when *n* = *1* (Fig. 1) because the infinite power series 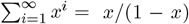 for |*x*| *<* 1. Incorporating this infinite power series into eqn. 3 shows that the expected proportion of predators handling prey at any given time will be *ahN* under the Type I. Importantly, this proportion differs from the expectation of zero that would be inferred to emerge by letting *h !* 0 in the way the Type I is typically derived (e.g., Holling, 1965; Rohr *et al*., 2022). In other words, the multi-prey model shows that handling times need not be inconsequential for the functional response to exhibit linear density dependence (Jeschke *et al*., 2004). Rather, even the Type I can be a very good approximation of feeding rates when *n* is high and less than 100% of predators are handling prey (i.e. *ahN <* 1), which requires that prey abundances remain less than 1*/ah*. For comparison, note that under the Type II the quantity 1*/ah* reflects the prey abundance at which 50% of predators will be handling prey (i.e. the per predator feeding rate is at half its maximum of 1*/h*), which is equivalent to the half-saturation constant of the Michaelis-Menten formulation. Of futher note is that under the multi-prey model 1*/ah* is also the prey abundance at which the proportions of predators handling 1, 2,. .., *n* prey are all equal (Fig. S.1).

### Empirical support for multi-prey feeding

The multi-prey model shows that a spectrum of functional response forms can exist between the extremes of the Type I and Type II when handling and searching are not assumed to be mutually exclusive (Fig. 1). This motivated us to test two main hypotheses using the large number of empirical functional response studies that exist in the literature. The first hypothesis was that prior syntheses indicating the Type I response to be rare (Dunn & Hovel, 2020; Jeschke *et al*., 2004) were biased against the Type I despite its potential empirical appropriateness. That is, feeding rates may have had response shapes between the Type II and rectilinear model (close to the Type I for prey abundances *<* 1*/ah*) but were classified as Type II due to the lack of a sufficiently simple rectilinear-approaching model in prior analyses. The second hypothesis was due to Sjöberg (1980) who motivated parameter *n* by considering it to be a measure of food particle size relative to a zooplankter’s gut capacity, with low *n* reflecting capacity for few large prey and high *n* reflecting capacity for many small prey. We thus expected predator-prey pairs with larger body-mass ratios to exhibit larger estimates of *n* when their functional responses were assumed to follow the multi-prey model. For generality and to safeguard against potential statistical model-comparison issues (see below), we included the Type I, II, III, multi-prey, and the generalized (eqn. 6) model in our comparisons. We were thus also able to test an additional hypothesis, due to Hassell *et al*. (1977), that larger body-mass ratios are associated with more pronounced Type III responses (i.e. larger values of *ϕ*).

We used the FoRAGE database of published functional response datasets to assess these hypotheses (Uiterwaal *et al*., 2022). Our v4 update contains 3013 different datasets representing 1015 unique consumer-resource pairs (i.e. not just predator and prey species, though we continue to refer to them as such for simplicity). For our analyses, we excluded datasets having a sample size less than 15 observations as well as structured experimental studies that implemented less than 4 different treatment levels of prey abundance (see the *Supplemental Materials* for additional details). Our model-fitting procedure followed the approach used by Stouffer & Novak (2021) and Novak & Stouffer (2021b), assuming one of two statistical models for each dataset: a Poisson likelihood for observational (field) studies and when eaten prey were replaced during the course of the experiment, and a binomial likelihood when eaten prey were not replaced. Experimental data available in the form of treatment-specific means and uncertainties were analyzed by a parametric bootstrapping procedure in which new datasets were created assuming either a treatment-specific Poisson or binomial process as dictated by the study’s replacement of prey. In cases where measures of the uncertainty around non-zero means were not available, we interpolated them based on the global log-log-linear relationship between means and standard errors across all datasets following Uiterwaal *et al*. (2018); for zero means, we interpolated missing uncertainty values assuming a linear within-dataset relationship. Unlike in Stouffer & Novak (2021) and Novak & Stouffer (2021b), we added a penalty to the likelihoods to discourage exceptionally large estimates of *n* and *ϕ* (see the *Supplementary Materials*) and bootstrapped data available in non-summarized form as well, using a non-parametric resampling procedure that maintained within-treatment sample sizes for treatment-structured datasets. Both replacement and non-replacement data were bootstrapped 100 times which was enough to obtain sufficient precision on the parameter point estimates.

### Frequency of multi-prey feeding

We used the Bayesian Information Criterion (BIC) to test our first hypothesis, counting the number of datasets whose bootstrapped mean BIC score supported a given model over the other models by more than two units (ΔBIC *>* 2). Our choice to use BIC was motivated both by its purpose of selecting the generative model (rather than the best out-of-sample predictive model, as per AIC) and by its generally stronger penalization of parametrically-complex models (thereby favoring simpler models, relative to AIC). Conclusions regarding evidence in support of the multi-prey model were thereby made more conservative, with our inclusion of models having equal or greater parametric complexity helping to guard against an inappropriate reliance on the asymptotic nature of BIC’s consistency property.

The result of this first analysis was that, overall, 925 (36%) of all 2,591 datasets provided support for functional response linearity (i.e. the Type I and multi-prey models), with 998 (38%) of all datasets providing support for multi-prey feeding more generally (i.e. the Type I, multi-prey, and generalized eqn. 6 models). When considering only those datasets that could differentiate among all five of the models, 7 (5.3%) of 132 replacement datasets and 143 (9.1%) of 1575 non-replacement datasets identified the multi-prey model (eqn. 5) as the sole best-performing model (Fig. 2a-2b). An additional 37 (28%) replacement and 451 (29%) non-replacement datasets identified the multi-prey model as performing equivalently well to their best-ranked model(s).

**Figure 2:**
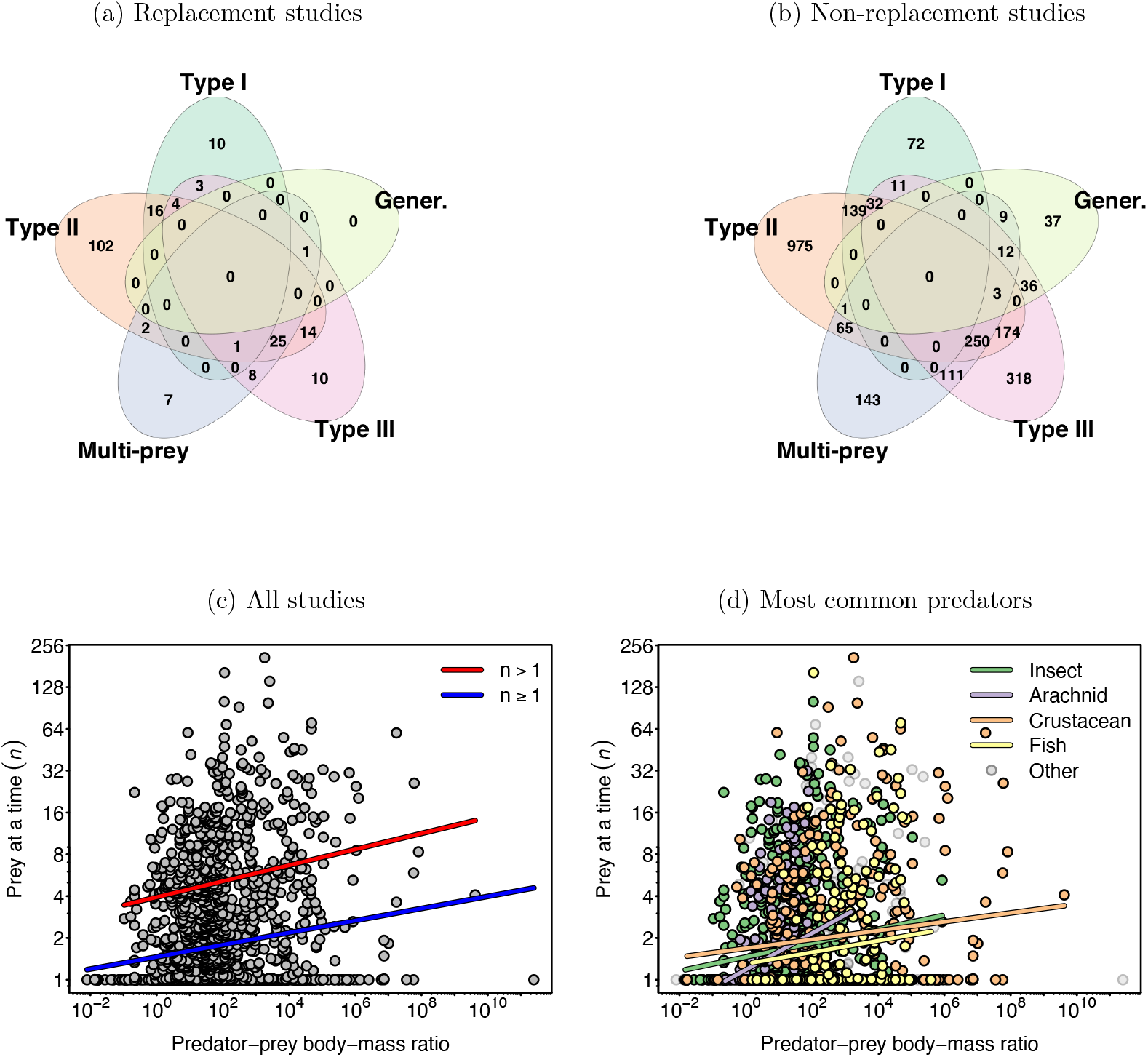
Empirical support for multi-prey feeding. Figs. 2a and 2b depict Venn diagrams categorizing the datasets of FoRAGE by their support for one or more of the five models as evaluated using a cut-off of 2 BIC units. Figs. 2c and 2d depict the observed relationship between estimates of *n* and the body-mass ratio of the studies’ predator-prey pairs, excluding datasets for which the Type I model alone performed best. Regression lines in Fig. 2c reflect all considered datasets or only those with estimates of *n >* 1 (Table S.1). Regression lines in Fig. 2d reflect the identity of the four most common predator groups (*n ≥*1, Table S.4).

Although the Type I and the generalized model were the least frequently sole-supported models, they were supported by datasets representing all four of the most common predator taxonomic groups that constituted 90% of all datasets in FoRAGE (insects, arachnids, crustaceans, and fishes; Fig. S.2).

### Effects of predator-prey body-mass ratio on *n* and *ϕ*

To test the second and third hypotheses, we excluded datasets for which the Type I had alone performed best and regressed the remaining datasets’ bootstrapped median point estimates of *n* and *ϕ* against their study’s predator-prey body-mass ratio (*ppmr*), these having been compiled in FoRAGE for most datasets. Although roughly 90% of these datasets had estimates of *n ≤* 8 and *ϕ ≤* 2 (Figs. S.3 and S.4), all three variables exhibited substantial variation in magnitude. We therefore performed linear least-squares regression using log_2_(*n*) and log_2_(*ϕ*) versus log_10_(*ppmr*). Our analysis supported the hypothesis that predator-prey pairs with larger body-mass ratios tend to exhibit larger estimates of *n* (Fig. 2c; *log*_2_(*n*) = 0.55 + 0.15 *· log*_10_(*ppmr*), *p <* 0.01, Table S.1), but the predictive utility of this relationship was extremely poor (*R*^2^ = 0.02). We also found support for the hypothesis that larger body-mass ratios are associated with larger values of *ϕ*, although the magnitude of this effect was weaker than it was for *n* (Fig. S.5; *log*_2_(*ϕ*) = 0.26 + 0.06 *· log*_10_(*ppmr*), *p <* 0.01, Table S.2) and was of similarly poor predictive utility (*R*^2^ = 0.02).

To assess the sensitivity of our result for *n* to variation among datasets, we performed additional regressions that restricted the considered datasets to (i) those having estimates of *n>* 1 (Fig. 2c, Table S.1), (ii) those with sample sizes exceeding the median sample size of all datasets (Fig. S.6, Table S.3), and (iii) the four most common predator taxonomic groups (insects, arachnids, crustaceans, and fishes), including for this last regression a two-way interaction term between predator group identity and predator-prey body-mass ratio (Fig. 2d, Table S.4). These analyses evidenced statistically clear, albeit predictively poor, positive relationships between *n* and predator-prey body-mass ratios for all predators in general and for each predator group individually as well.

### Population-dynamic effects of multi-prey feeding

Given the empirical evidence that multi-prey feeding may indeed be common and a viable way to describe functional responses, we next investigated its potential consequences for predator-prey dynamics. Our goal was to understand how assuming either a Type I or Type II response could lead to incorrect conclusions regarding these dynamics. We used the well-studied Rosenzweig & MacArthur (1963) model to achieve this goal, employing graphical (i.e. isocline) analysis and both deterministic and stochastic simulations.

The model describes the growth rates of the prey *N* and predator *P* populations as

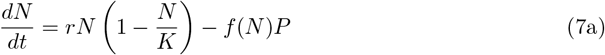

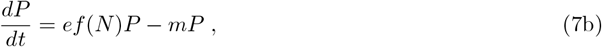

where *r* and *K* are the prey’s intrinsic growth rate and carrying capacity, *f* (*N*) is the functional response, and *e* and *m* are the predator’s conversion efficiency and mortality rate. Logistic prey growth and Holling’s Type II response have become the component parts of the canonical Rosenzweig-MacArthur model for which enrichment in the form of an increasing carrying capacity causes the populations’ dynamics to transition from a regime of monotonically-damped stable coexistence to damped oscillations to sustained limit cycles (Rosenzweig, 1971). Other prey growth and Type II-like functional response forms affect a similar destabilization sequence (e.g., Freedman, 1976; May, 1972; Rosenzweig, 1971; Seo & Wolkowicz, 2018). The location of the Hopf bifurcation between asymptotic stability and limit cycles is visually discerned in the model’s *P* vs. *N* phase plane (Fig. 3) as the point where the vertical *N* ^*^ predator isocline intersects the parabolic *P* ^*^ prey isocline at its maximum, half-way between *-*1*/ah* and *K* (Rosenzweig, 1969; Rosenzweig & MacArthur, 1963). That is, the coexistence steady state entails a globally-stable fixed point when the isoclines intersect to the right of the maximum and entails a locally-unstable fixed point with a globally-stable limit cycle when they intersect to the left of the maximum (Seo & Wolkowicz, 2018). Graphically, increasing *K* destabilizes dynamics by stretching the prey isocline, moving its maximum to the right while the position of the vertical predator isocline remains unchanged. In contrast, when logistic growth and a Type I are assumed, the prey isocline is a linearly-decreasing function of prey abundance (Fig. 3) and predator-prey coexistence entails a globally-stable fixed point for all levels of enrichment.

**Figure 3:**
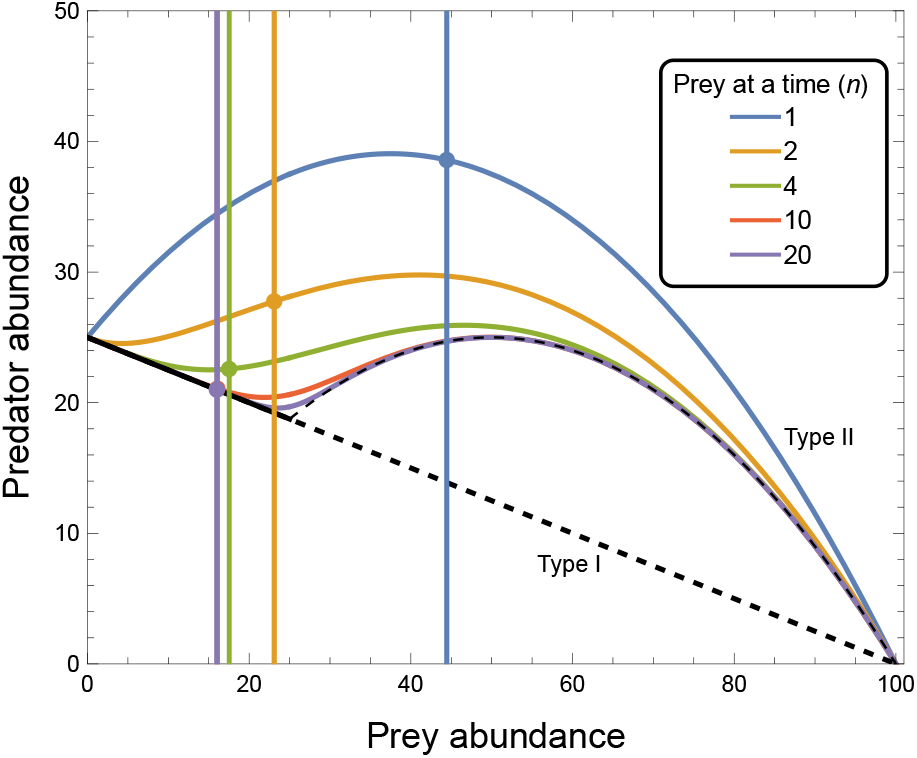
Predator and prey isoclines of the Rosenzweig-MacArthur model modified to include the multi-prey response correspond to those observed with the Type I and Type II responses when *n* = *1* and *n* = 1 respectively. As the number *n* of prey that a predator can handling while searching increases, the prey abundance at which the predator’s growth rate is zero (i.e. the vertical predator isocline, *N* ^*^) decreases from its value under the Type II response (*m/a*(*e-mh*)) and converges rapidly on the value expected under the Type I response (*m/ae*). In contrast, predator abundances at which the prey’s growth rate is zero, *P* ^*^, converge on those expected under the Type I response only at low prey abundances to affect a second region of asymptotically stable dynamics; the “hump” does not flatten as it would if the handling time were presumed to be inconsequential (i.e. *h* = 0). Limit cycles occur when the predator and prey isoclines intersect on the left flank of the hump. With increasing *n*, the inflection point between the low-prey region of stability and the intermediate region of limit cycles approaches the prey abundance where all predators become busy handling prey under the rectilinear model, 1*/ah* (indicated by non-dashed region of the black prey isocline). *Other parameter values*: attack rate *a* = 0.02, handling time *h* = 2, prey growth rate *r* = 0.5, prey carrying capacity *K* = 100, conversion efficiency *e* = 0.25, predator mortality rate *m* = 0.08.

### Graphical analysis

For our analysis we insert the multi-prey response (eqn. 5) for *f* (*N*) in eqn. 7. Solving *dP/dt* = 0 for the *N* ^*^ predator isocline then requires solving

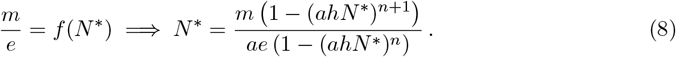

This leads to a solution for *N* ^*^ that is independent of the predator’s abundance (i.e. remains vertical in the *P* vs. *N* phase plane) but is unwieldy for *n >* 2 (see *Supplementary Materials*).

Nonetheless, it represents a generalization of the predator isocline obtained for the Rosenzweig-MacArthur model with 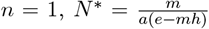,and converges on *N* ^*^ = *m/ae* as *n* when *ahN* ^*^ *<* 1, just as obtained assuming the Type I. In fact, *N* ^*^ transitions smoothly from the former to the latter as *n* increases (Fig. 3) because eqn. 8 is a monotonically declining function of *n* for *ahN* ^*^ *<* 1.

Solving *dN/dt* = 0 for the *P* ^*^ prey isocline leads to the solution

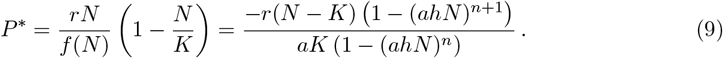

This too represents a generalization of the Rosenzweig-MacArthur model’s prey isocline, *P* ^*^ = *-*(*r/aK*)(*N -K*)(1+*ahN*), which is itself a generalization of the isocline *P* ^*^ = *-*(*r/aK*)(*N -K*) obtained with the Type I as *n ! 1*. Between these the prey isocline under the multi-prey response transitions from a parabolic dependence on the prey’s abundance to having a second region within which it is a declining function of prey abundance (Fig. 3). This second region has a slope of *-*(*r/aK*) at its origin regardless of *n* and is limited to low prey abundances of *N <* 1*/ah*; as *n* increases, the region’s upper extent approaches the prey abundance at which all predators are busy handling prey under the rectilinear model. That is, for 1 *< n < 1* the “hump” shape of *P* ^*^ does not flatten out as it does when one assumes handling times to become negligible. Rather, the *P* ^*^ converges on *-*(*rhN/K*)(*N - K*) for *N >* 1*/ah* as *n* increases and thus, similar to what can occur for the Type III response (Uszko *et al*., 2015), exhibits two regions of negative prey dependence (where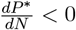) that flank an intermediate region of positive prey dependence (where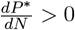).

### Implications for coexistence and dynamics

The emergence of a second prey abundance region where the slope of the prey isocline is negative means that a second asymptotically-stable coexistence equilibrium — one having a high predator-to-prey abundance ratio — is possible should the two isoclines intersect within it. The fact that this may occur is discerned by noting that *N* ^*^ (eqn. 8) is independent of *r* and *K*, and that *P* ^*^ (eqn. 9) is independent of *m* and *e*; the positions of the two isoclines are thus independent except via the functional response parameters *a, h*, and *n*. In fact, because *N* ^*^ decreases while the upper limit of the low prey abundance region of *P* ^*^ increases towards 1*/ah* as *n* increases, it is readily possible — conditional on the values of the other parameters — to observe a stable state at *n* = 1 to first transition to limit cycles and then return to fixed-point stability as *n* alone is increased. This is illustrated by Fig. 4 in the context of enrichment for values of *K* between approximately 75 and 115. Multi-prey feeding may thus be seen as another mechanism contributing to stability at high productivity (Roy & Chattopadhyay, 2007). Indeed, in addition to rescuing predators from deterministic extinction at low levels of enrichment where a single-prey-at-a-time predator could not persist (20 *< K <* 40 in Fig. 4), sufficiently large values of *n* can preclude the occurrence of limit cycles altogether (*n>* 9 in Fig. 4).

**Figure 4:**
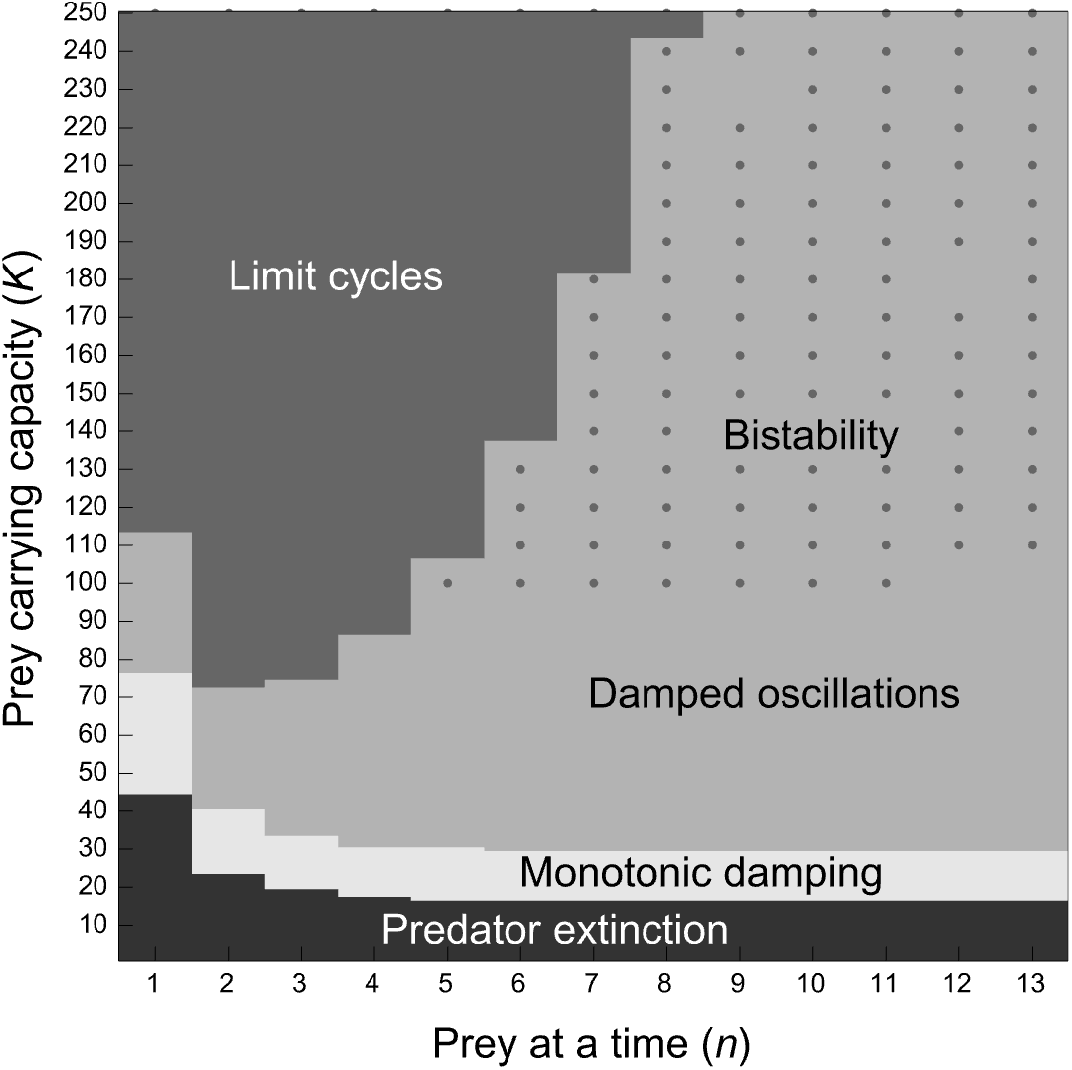
The destabilization with enrichment that is seen under the classic Rosenzweig-MacArthur model (where *n* = 1) is altered when predators can search for and handle multiple prey at a time (*n >* 1). At low prey carrying capacities (*K <* 40), multi-prey feeding rescues predators from deterministic extinction. At intermediate carrying capacities (40 *< K <* 110), low levels of multi-prey feeding destabilize dynamics by causing perturbation responses to transition from a transient regime of monotonic damping to one of damped oscillations or from damped oscillations to a persistent limit cycle regime. Further increases in multi-prey feeding can have a qualitatively stabilizing influence on dynamics, with sufficiently high *n* precluding a transition to limit cycles altogether so long as perturbations are sufficiently small. Large perturbations, on the other hand, will cause a transition to an alternative stable state consisting of limit cycle dynamics (see Fig. 5). *Other parameter values* as in Fig. 3.

Notably, however, the just-described high-predator low-prey steady state is only a locally stable fixed point and coexists with a stable limit cycle that surrounds it (Figs. 4 and 5). The high-predator low-prey state thus exhibits bi-stability. The consequences of this bi-stability are that predator-prey interactions with multi-prey feeding are destined to exhibit (i) transitions to persistent limit cycles when subjected to large perturbations that send abundances beyond the domain of attraction of the fixed-point steady state (Fig. 5*a,c*), and (ii) transient dynamics that are prone to damped oscillations (rather than monotonic damping) in response to small perturbations within the domain of attraction. These transient oscillations occur for substantially lower levels of enrichment than is the case for single prey-at-a-time predators (Fig. 4). Moreover, their temporal duration can be exceedingly long (Fig. 5*b*) because the limit cycle acts akin to a crawl-by attractor (Hastings *et al*., 2018) that impinges upon the steady state’s local resilience. Thus, when subjected to continual perturbations in an explicitly stochastic setting (Barraquand *et al*., 2017), the system can readily transition between the stable fixed-point attractor and the stable limit cycle attractor that surrounds it (Fig. 6), resulting in dynamical epochs of irregular duration that are characteristic of many empirical time-series (Blasius *et al*., 2020; Rubin et al., 2023). Therefore, multi-prey feeding does not provide a robust mechanism against instability at high productivity but rather leads to a richer range of population dynamics and coexistence states than can result from Type I, II, or III responses alone.

**Figure 5:**
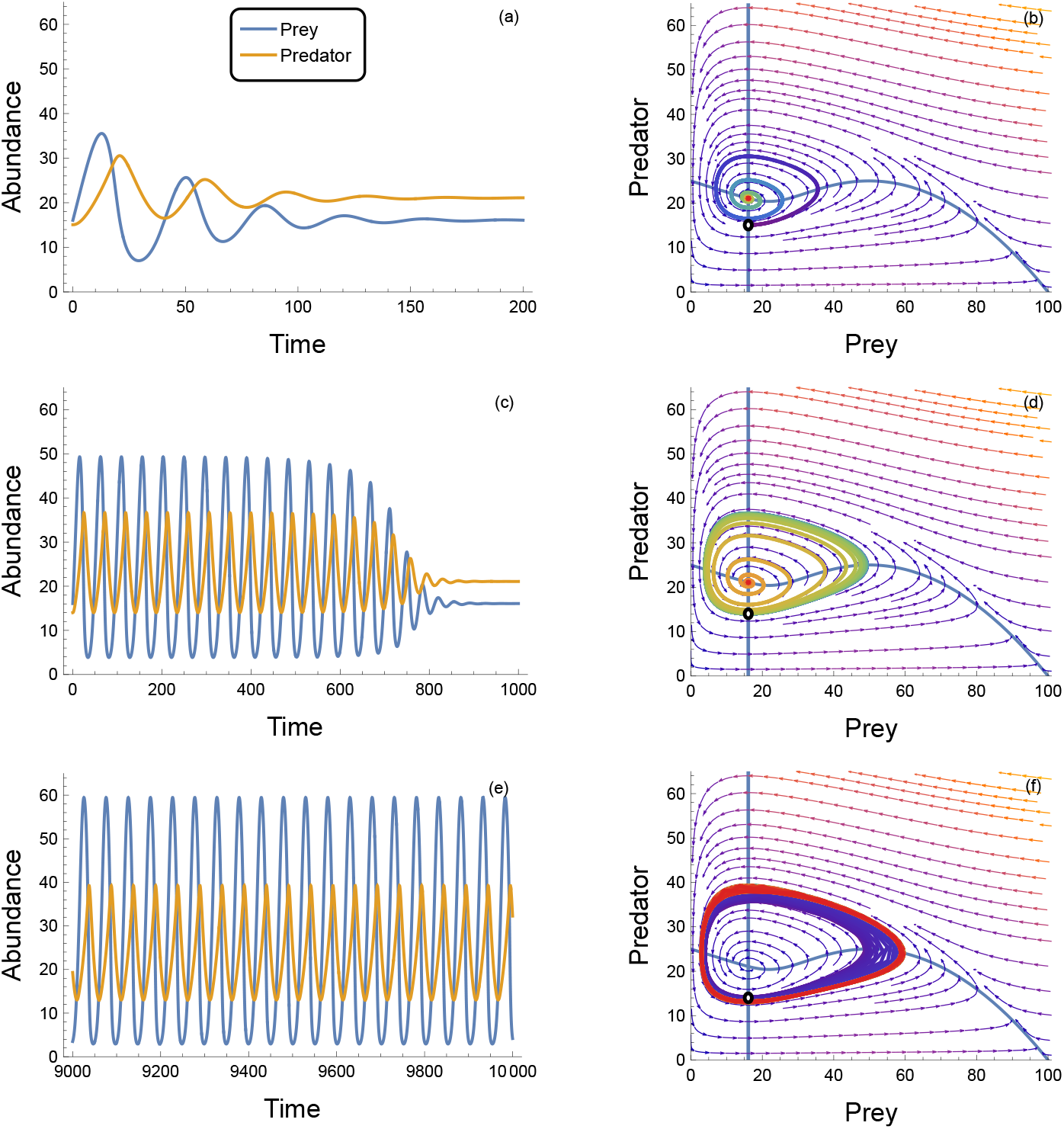
Because of the system’s bi-stability at high predator-to-prey abundance ratios, even small differences in the size of a perturbation to the steady state can affect a large change in the duration of the system’s transient response (compare panels *a* and *b* with *c* and *d*) and can even cause the system to become entrained in a stable limit cycle (illustrated in panels *e* and *f*). The only difference between each of the above panel rows is that the predator’s initial population size *P* (0) is perturbed away from its *P* ^*^ steady state as: (*a, b*) *P* (0) = *P* ^*^ − 6; (*c, d*) *P* (0) = *P* ^*^ − 7.0645; and (*e, f*) *P* (0) = *P* ^*^ − 7.065. For all cases *N* (0) = *N* ^*^. *Parameter values* as in Fig. 3 with *n* = 10.

**Figure 6:**
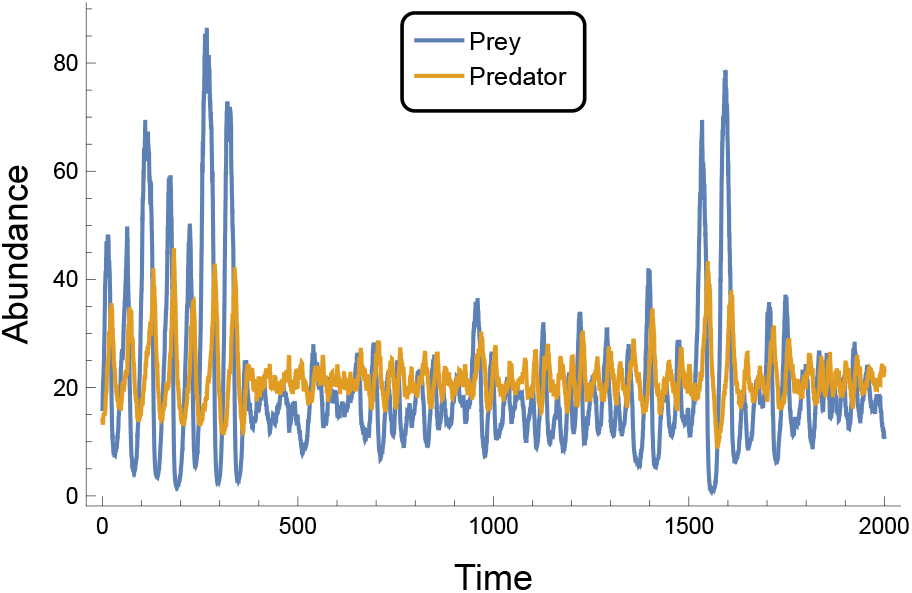
When subjected to continually-occurring stochastic perturbations, the high-predator low-prey coexistence state can exhibit time periods during which its dynamics are influenced primarily by the stable fixed-point attractor and time periods during which dynamics are primarily influenced by the alternative stable limit cycle attractor, switching between these on an irregular basis. Simulation implemented using Itô stochastic differential equations as *dN* = *rN* (1 − *N/K*) − *f* (*N*)*P dt* + *σNdW* and *dP* = *ef* (*N*) − *mP dt* − *σPdW*, with *f* (*N*) as in eqn. 5 and Gaussian white environmental noise *dW* (*t*) of volatility *σ*= 0.04 (*cf*. Barraquand, 2023). *Other parameter values* and initial population sizes as in Fig. 5c-d.

## Discussion

Our study was motivated by the apparent disconnect that exists between the way that many empirically-minded ecologists perceive functional response linearity and the way that many modelers and theory-minded ecologists justify its use in their representations of consumer-resource interactions. While the former are prone to dismiss the Type I as being overly simplistic and hence unsuitable for describing predator feeding rates, the latter are prone to rely on and justify its sufficiency for the sake of computational ease and analytically-tractable insight. Since the potential for predators to feed on multiple prey at a time (i.e. the non-exclusivity of handling and searching activities) has been little considered by either group, we set out to address three aspects of this disconnect: (i) deriving a multiple-prey-at-a-time model that mechanistically connects the linear and rectilinear models to the more empirically palatable Type II model, (ii) assessing the extent to which published datasets provide support for multi-prey feeding, and (iii) investigating how multi-prey feeding and the linear density dependence it can impose on feeding rates can alter our understanding of predator-prey coexistence. Because they bear insight with which to elaborate on the circumstances under which linearity may be empirically relevant, we structure the discussion of our work by considering the latter two aspects first.

### Empirical support

Our statistical analysis of the datasets compiled in FoRAGE demonstrates that both the Type I and multi-prey models are viable descriptions (*sensu* Skalski & Gilliam, 2001) of the feeding rates that predators have exhibited in many single-prey experiments (Figs. 2a-2b). This result is consistent with handling and searching being non-exclusive activities for a substantial number of predator-prey pairs. Although our result contrasts with the prior syntheses of Jeschke *et al*. (2004) and Dunn & Hovel (2020), these (*i*) did not consider models capable of response forms in between the strictly linear Type I and Type II forms and (*ii*) either relied on the conclusions reached by each studies’ original authors (who used varied model-fitting and comparison approaches) or visually assessed functional response forms from plotted data. One might argue that many of the datasets providing sole support to the Type I in our analysis came from experiments using prey abundances that were insufficient to elicit saturation (see also Coblentz *et al*., 2023), but the point can be made that, from an information-theoretic perspective, the Type I performed best across the range of prey abundances that the original authors considered empirically reasonable (and logistically feasible). The even greater number of datasets that provided sole support to the multi-prey model, along with the result that many of the point estimates for parameter *n* (the maximum number of prey eaten at a time) were sufficiently large to affect a response approaching a rectilinear response (Figs. 1 and 2c), indicates that feeding rates exhibited a region of linearity for many predator-prey interactions having long handling times as well. Moreover, the statistically-clear positive relationships we observed in our subsequent regression analyses of *n* and predator-prey body-mass ratios (Figs. 2c-2d) support Sjöberg’s hypothesis regarding a proximate reason for this linearity, it being more likely to occur for larger predators feeding on small prey because handling is less preclusive of searching.

Unfortunately, the amount of variation in *n* that was explained by body-mass ratio alone was extremely low, making the relationship of little predictive utility relative to several other body-mass relationships (e.g., Brose *et al*., 2006; Coblentz et al., 2023; Hatton et al., 2015; Rall et al., 2012). That said, the relationship’s low explanatory power is not unsurprising given that none of the experiments in FoRAGE was designed with the multi-prey model in mind. In particular, and although most estimates of *n* were of a seemingly reasonable magnitude (Fig. S.3), we caution against giving too much credence to the very large-valued estimates we observed. This is for two primary reasons. First, given that a given dataset’s ability to distinguish between possible values of *n* diminishes rapidly as *n* increases (Fig. 1), datasets exhibiting saturation at high prey abundances but having few or no observations near the inflection point of 1*/ah* will have been sensitive to issues of parameter identifiability. Low identifiability will have caused an inflation of estimates despite our effort to guard against it by removing datasets with fewer than 4 prey abundance levels. Second, given that initiating experiments with predator individuals having empty guts is a common protocol (Griffen, 2021; Li *et al*., 2018), many experiments will have strictly violated the assumption of predator behavior being at steady state. This will also have inflated estimates of *n* by causing transient rates of prey ingestion to exceed rates of handling completion (i.e. *aN >* 1*/h*) to affect faster-than-steady-state feeding, especially at prey abundances below 1*/ah*. We therefore suggest that the very large estimates of *n* observed in our analyses be better interpreted as qualitative (rather than quantitative) support for the non-exclusivity of searching and handling and encourage future experiments and analyses with additional covariate predictors to better understand the biological sources of variation in *n*. (Similar issues pertain to the estimation and interpretation of *ϕ*.)

### Mechanistic approximations

The multi-prey model may be considered a mechanistic model in that its derivation and each of its parameters has at least one biologically-specific interpretation. However, it is also rather phenomenological in that it encodes only an essence of the biologically possible non-exclusivity of searching and handling processes. For example, the model’s derivation assumes that the attack rate and handling time remain constant and independent of the number of prey that predators are already handling (below the maximum number *n*). Although this assumption may result in a very good approximation to feeding rates, it is unlikely to reflect biological reality particularly as the number of prey being handled by a given predator approaches *n*. In such circumstances either or both searching and handling process rates are likely to become dependent on the feeding rate and thereby on prey abundance (see also Okuyama, 2010; Stouffer & Novak, 2021).

Functional responses where such dependence is important may be better and more mechanistically described by more flexible models (see also Novak & Stouffer, 2021a). Prominent among these is the extended Steady State Saturation model (SSS^1^) of Jeschke *et al*. (2004) in which handling and digestion are explicitly distinguished (see *Supplementary Materials*). In this four-parameter model, searching and handling are mutually exclusive, but searching and digestion are not because the predator’s search effort depends on its gut fullness (i.e. hunger level) and is thus dictated by the digestion rate. A phenomenological shape parameter controls the non-linearity of the search-effort hunger-level relationship. For high values of this shape parameter (reflecting predators that search at their maximum rate even when their guts are quite full) and inconsequential handling times, the model approaches the rectilinear model, just like the multi-prey model at high *n*, while for consequential handling times it retains a saturating curvature at low prey abundances (see Figs. A1 and A2 of Jeschke *et al*., 2004).

### Population-dynamic effects

The population-dynamic consequences of the extended SSS model remain unstudied, but our analysis of the simpler multi-prey model reveals the relevance of it and other models for understanding how the linearity of multi-prey feeding can impact predator-prey dynamics. These other models are the arctangent and hyperbolic tangent models because for these it has been more rigorously shown that two limit cycles — one stable and the other unstable — can co-occur with a locally-stable fixed point at low prey abundances (Seo & Kot, 2008; Seo & Wolkowicz, 2015; 2018), just as we observed for the multi-prey model (see also Freedman, 1980). The key feature common to all three models is that they affect a prey isocline that *decreases* from a *finite*-valued origin at zero prey abundance. This differs from the Type II and other functional responses that are concave and increasing with prey density at low prey abundance. For these the prey isocline *increases* from a finite-valued origin, the low-prey fixed point is unstable, and only the stable limit cycle is thus of relevance under logistic prey growth. It also differs from functional responses that accelerate at low prey abundances (e.g., the Type III) and from consumer-resource models more generally in which, for example, prey have a physical refuge, exhibit sublinear density-dependence, or experience density-independent immigration. For these the prey isocline decreases from an origin that approaches infinity and the low prey steady state is a stable fixed point around which limit cycles do not occur (e.g., Case, 2000; Uszko *et al*., 2015). We surmise that the linearity brought about by the non-exclusivity of searching and handling in the multi-prey model is (*i*) replicated by the more phenomenological arctangent and hyperbolic tangent models, and that (*ii*) it is the cause of the greater range of dynamical outcomes that these functional responses affect as compared to responses exhibiting nonlinearity at low prey abundances.

The broader implication of the multi-prey model is that the conclusions and predictions of simple consumer-resource theory which relies on the linear Type I may not be as broadly predictive of population and ecosystem dynamics as the mathematics would suggest. More specifically, the multi-prey model shows that such theory’s domain of relevance to natural systems, in which consumers invariably have a (potentially unobserved) maximum feeding rate, is limited to quantifiably small perturbations. Our consideration of enrichment effects illustrates an example of this. If a focal predator’s functional response were assumed to be linear Type I, species’ fixed point abundances would be inferred to be globally stable, with perturbations decaying monotonically regardless of the enrichment level. In contrast, if the predator were to be correctly recognized as being able to feed on multiple prey at a time even as its functional response appeared linear based on observations or experiments, then the same fixed point abundances would be recognized as being only locally stable, with sufficiently large perturbations predicted to elicit cycles that could persist for many generations or even indefinitely. Indeed, as indicated by Rubin *et al*. (2023) in their analysis of a stochastic implementation of the Rosenzweig-MacArthur model, the real-world dynamics would additionally be influenced by the crawl-by inducing origin (dual extinction) and prey-only (carrying capacity) steady states that can extend the lifetime of long-term transients even further. The influence of these phenomena, too, would not be inferred to be important were a linear Type I to be assumed because these unstable steady states would rarely if ever be approached during simulation forecasts.

### Relevance revisited

As discussed above (see *Relevance of Type I response*), the multi-prey model shows that handling times need not be inconsequential to observe linear prey dependence when the number of prey that a predator individual can handle at a time is relatively high and the maximum proportion of individuals in a predator population that are simultaneously handling prey remains sufficiently low. This is not to say that other factors and processes cannot cause functional responses to be very nonlinear, but within the confines of our work’s assumptions the latter condition can be satisfied as long as prey abundances remain less than 1*/ah*.

Our statistical and mathematical analyses add insight into when the conditions for linearity are more likely to be met. Specifically, functional responses are more likely to exhibit linearity when predator-to-prey body-mass ratios are high (Fig. 2c), when predator-to-prey abundance ratios are high (Fig. 3), and thus, we predict, in top-heavy systems with high predator-to-prey biomass ratios. Top-heavy interactions and food webs more generally occur in all ecosystem types (McCauley *et al*., 2018), but are more likely for ectothermic and invertebrate consumers, in aquatic habitats, among higher trophic levels, and in ecosystems of low total biomass (Brose *et al*., 2006; Hatton *et al*., 2015; Perkins *et al*., 2022). The development of methods for gauging the nonlinearity of functional responses in diverse field settings (e.g., Novak *et al*., 2017; Uiterwaal & DeLong, 2024) will be useful for directly testing our prediction that these same systems should also exhibit more linear functional responses. New methods that make use of the greater information content associated with counts of the numbers of prey being handled (Fig. S.1) should be particularly useful.

Importantly, our work also shows that predator-prey dynamics need not be destabilized by food web top-heaviness. Rather, paralleling theory assuming Type III responses (Kalinkat *et al*., 2013; Uszko et al., 2015), increases in top-heaviness can lead to greater food web stability — be it stable coexistence potential or perturbation resilience (Fig. S.7) — when multi-prey feeding occurs, provided that perturbations are small enough for population abundances to remain well within the local attractor of the stable fixed point (Fig. 5). This contrasts with existing theory on top-heavy food webs that has largely assumed Type II responses (McCauley *et al*., 2018). Indeed, our analyses show that even small departures from mutual exclusivity can lead to qualitatively different coexistence states and dynamics than predicted by existing theory, including the possibility of long-term transients and the just-mentioned bi-stability of fixed-point and limit-cycle dynamics. Food web models that incorporate multi-prey feeding and how its prevalence may change with species- and system-level attributes will be useful for understanding just how much multi-prey feeding must occur within food webs as a whole to alter community structure and dynamics. A first step towards such food web models will be to extend the multi-prey model to multi-species formulations appropriate for generalist rather than single-prey-species predators.

### Conclusions for bridging theory and empirical insight

Natural history observations show that diverse types of predators are capable of (literally) handling and searching for prey simultaneously: sea otters capture several snails on a dive; crabs process mussels with their mouthparts while picking up more with their claws; spiders capture insects in their webs while processing others for later ingestion. Many more situations relevant to multi-prey feeding become apparent and potentially relevant to the context of functional responses when it is recognized that the “handling time” parameter of most models represents not just the literal manipulation of prey (e.g., that may be seen by an observer of the interaction) but rather reflects the feeding process that limits a predator’s maximum feeding rate, including possible limits to stomach fullness and digestion (DeLong, 2021; Jeschke *et al*., 2002; 2004). Sculpin fishes, for example, have been observed with over 300 identifiable mayflies in their stomachs (Preston *et al*., 2018), the majority of which could not have been captured simultaneously and for which literal handling must therefore have been inconsequential relative to digestion.

The degree to which searching and (general) handling actually represent mutually exclusive activities, and the degree to which each of the many processes potentially encapsulated by a handling time parameter measurably contributes to a predator’s functional response, is nonetheless poorly discerned from observation alone. Knowing that handling times are short or long, or that searching and literal handling do or do not overlap, is neither sufficient to dismiss or assume a given functional response model on *a priori* grounds. This is because all models are phenomenological approximations of biological process at some level. This applies as much to predator-prey interactions studied in controlled experiments as it does to those studied in natural settings, and is particularly true in the context of building understanding and theory when extrapolating the former to the latter across Ecology’s wide-ranging scales. In this context we draw two overarching conclusions from our analyses: that functional response linearity should not be dismissed by empiricists as an irrelevant description of predator feeding rates, and that modelers and theoreticians should be more cautious in reaching empirical conclusions of system dynamics when presuming the linear Type I response to be appropriate.

## Supporting information

SOM

## Acknowledgments

MN thanks the OSU MathBio group for feedback, is indebted to Patrick DeLeenheer for setting him straight, and thanks CJ Keist for technical assistance with OSU’s Cosine High Performance Cluster. We also thank Frédéric Barraquand, Wojciech Uszko, and Matthieu Barbier for helping us improve the manuscript. A preprint version of this article has been peer-reviewed and recommended by PCIEcology (https://doi.org/10.24072/pci.ecology.100702).

## Supplementary Online Information

I. Multi-prey functional response model; II. Analysis of FoRAGE datasets; III. Population-dynamic effects; IV. A reformulation of the extended Steady State Saturation model.

## Footnotes

1 We would be remiss not to point out that all functional response models of which we are aware assume steady state conditions at the behavioral foraging scale. The SSS model’s name does not, therefore, reflect a limitation that is unique to it.

## References

Barraquand, F. (2023). No sensitivity to functional forms in the Rosenzweig-MacArthur model with strong environmental stochasticity. Journal of Theoretical Biology, 572, 111566.

Barraquand, F., Louca, S., Abbott, K. C., Cobbold, C. A., Cordoleani, F., DeAngelis, D. L., Elderd, B. D., Fox, J. W., Greenwood, P., Hilker, F. M., Murray, D. L., Stieha, C. R., Taylor, R. A., Vitense, K., Wolkowicz, G. S. K. & Tyson, R. C. (2017). Moving forward in circles: challenges and opportunities in modelling population cycles. Ecology Letters, 20, 1074–1092.

Baudrot, V., Perasso, A., Fritsch, C., Giraudoux, P. & Raoul, F. (2016). The adaptation of generalist predators’ diet in a multi-prey context: insights from new functional responses. Ecology, 97, 1832–1841.

Blasius, B., Rudolf, L., Weithoff, G., Gaedke, U. & Fussmann, G. F. (2020). Long-term cyclic persistence in an experimental predator–prey system. Nature, 577, 226–230.

Brose, U., Jonsson, T., Berlow, E. L., Warren, P., Banasek-Richter, C., Bersier, L.-F., Blanchard, J. L., Brey, T., Carpenter, S. R., Blandenier, M.-F. C., Cushing, L., Dawah, H. A., Dell, T., Edwards, F., Harper-Smith, S., Jacob, U., Ledger, M. E., Martinez, N. D., Memmott, J., Mintenbeck, K., Pinnegar, J. K., Rall, B. C., Rayner, T. S., Reuman, D. C., Ruess, L., Ulrich, W., Williams, R. J., Woodward, G. & Cohen, J. E. (2006). Consumer-resource body-size relationships in natural food webs. Ecology, 87, 2411–2417.

Case, T. J. (2000). Illustrated Guide to Theoretical Ecology. Oxford University Press, New York.

Coblentz, K. E., Novak, M. & DeLong, J. P. (2023). Predator feeding rates may often be unsaturated under typical prey densities. Ecology Letters, 26, 302–312.

DeLong, J. P. (2021). Predator ecology: Evolutionary ecology of the functional response. Oxford University Press.

DeLong, J. P. & Uiterwaal, S. F. (2018). The FoRAGE (Functional Responses from Around the Globe in all Ecosystems) database: a compilation of functional responses for consumers and parasitoids. Knowledge Network for Biocomplexity, doi:10.5063/F17H1GTQ.

Denny, M. (2014). Buzz Holling and the functional response. The Bulletin of the Ecological Society of America, 95, 200–203.

Dunn, R. P. & Hovel, K. A. (2020). Predator type influences the frequency of functional responses to prey in marine habitats. Biology letters, 16, 20190758.

Freedman, H. (1976). Graphical stability, enrichment, and pest control by a natural enemy. Mathematical Biosciences, 31, 207–225.

Freedman, H. I. (1980). Deterministic mathematical models in population ecology. Pure and Applied Mathematics. Marcel Dekker, Inc., New York, NY, USA.

Garay, J. (2019). Technical review on derivation methods for behavior dependent functional responses. Community Ecology, 20, 28–44

Griffen, B. D. (2021). Considerations when applying the consumer functional response measured under artificial conditions. Frontiers in Ecology and Evolution, 9, 461.

Hassell, M. P., Lawton, J. H. & Beddington, J. R. (1977). Sigmoid functional responses by invertebrate predators and parasitoids. The Journal of Animal Ecology, 46, 249–262.

Hastings, A., Abbott, K. C., Cuddington, K., Francis, T., Gellner, G., Lai, Y.-C., Morozov, A., Petrovskii, S., Scranton, K. & Zeeman, M. L. (2018). Transient phenomena in ecology. Science, 361.

Hatton, I. A., McCann, K. S., Fryxell, J. M., Davies, T. J., Smerlak, M., Sinclair, A. R. E. & Loreau, M. (2015). The predator-prey power law: Biomass scaling across terrestrial and aquatic biomes. Science, 349, aac6284.

Holling, C. S. (1959a). The components of predation as revealed by a study of small-mammal predation of the european pine sawfly. The Canadian Entomologist, 91, 293–320.

Holling, C. S. (1959b). Some characteristics of simple types of predation and parasitism. The Canadian Entomologist, 91, 385–398.

Holling, C. S. (1965). The functional response of predators to prey density and its role in mimicry and population regulation. Memoirs of the Entomological Society of Canada, 45, 3–60.

Jeschke, J. M. (2007). When carnivores are “full and lazy”. Oecologia, 152, 357–364.

Jeschke, J. M., Kopp, M. & Tollrian, R. (2002). Predator functional responses: Discriminating between handling and digesting prey. Ecological Monographs, 72, 95–112.

Jeschke, J. M., Kopp, M. & Tollrian, R. (2004). Consumer-food systems: why type I functional responses are exclusive to filter feeders. Biological Reviews, 79, 337–349.

Kalinkat, G., Rall, B. C., Uiterwaal, S. F. & Uszko, W. (2023). Empirical evidence of type III functional responses and why it remains rare. Frontiers in Ecology and Evolution, 11.

Kalinkat, G., Schneider, F. D., Digel, C., Guill, C., Rall, B. C. & Brose, U. (2013). Body masses, functional responses and predator–prey stability. Ecology Letters, 16, 1126–1134.

Koen-Alonso, M. (2007). A process-oriented approach to the multispecies functional response. In: From Energetics to Ecosystems: The Dynamics and Structure of Ecological Systems (eds. Rooney, N., McCann, K. S. & Noakes, D. L. G.). Springer, Dordrecht, pp. 2–32.

Li, Y., Rall, B. C. & Kalinkat, G. (2018). Experimental duration and predator satiation levels systematically affect functional response parameters. Oikos, 127, 590–598.

Lotka, A. J. (1925). Elements of physical biology. Williams & Wilkins.

May, R. M. (1972). Limit cycles in predator-prey communities. Science, 177, 900–902.

McCauley, D. J., Gellner, G., Martinez, N. D., Williams, R. J., Sandin, S. A., Micheli, F., Mumby, P. J. & McCann, K. S. (2018). On the prevalence and dynamics of inverted trophic pyramids and otherwise top-heavy communities. Ecology Letters, 21, 439–454.

Mills, N. J. (1982). Satiation and the functional response: a test of a new model. Ecological Entomology, 7, 305–315.

Murdoch, W. W. & Oaten, A. (1975). Predation and population stability. Advances in Ecological Research, 9, 1–131.

Novak, M. (2010). Estimating interaction strengths in nature: experimental support for an observational approach. Ecology, 91, 2394–2405.

Novak, M., Coblentz, K. E. & DeLong, J. P. (2025a). Data: In defense of the original Type I functional response: The frequency and population-dynamic effects of feeding on multiple prey at a time. FigShare 10.6084/m9.figshare.28292147.v1.

Novak, M., Coblentz, K. E. & DeLong, J. P. (2025b). Supplementary Online Materials: In defense of the original Type I functional response: The frequency and population-dynamic effects of feeding on multiple prey at a time. FigShare 10.6084/m9.figshare.28292213.v1.

Novak, M. & Stouffer, D. B. (2021a). Geometric complexity and the information-theoretic comparison of functional-response models. Frontiers in Ecology and Evolution, 9, 776.

Novak, M. & Stouffer, D. B. (2021b). Systematic bias in studies of consumer functional responses. Ecology Letters, 24, 580–593.

Novak, M. & Stouffer, D. B. (2024). Corrigendum: Geometric complexity and the informationtheoretic comparison of functional-response models. Frontiers in Ecology and Evolution, 12.

Novak, M., Wolf, C., Coblentz, K. E. & Shepard, I. D. (2017). Quantifying predator dependence in the functional response of generalist predators. Ecology Letters, 20, 761–769.

Okuyama, T. (2010). Prey density-dependent handling time in a predator-prey model. Community Ecology, 11, 91–96.

Perkins, D. M., Hatton, I. A., Gauzens, B., Barnes, A. D., Ott, D., Rosenbaum, B., Vinagre, C. & Brose, U. (2022). Consistent predator-prey biomass scaling in complex food webs. Nature Communications, 13, 4990.

Preston, D. L., Henderson, J. S., Falke, L. P., Segui, L. M., Layden, T. J. & Novak, M. (2018). What drives interaction strengths in complex food webs? A test with feeding rates of a generalist stream predator. Ecology, 99, 1591–1601.

Rall, B. C., Brose, U., Hartvig, M., Kalinkat, G., Schwarzmüller, F., Vucic-Pestic, O. & Petchey, O. L. (2012). Universal temperature and body-mass scaling of feeding rates. Philosophical Transactions of the Royal Society B: Biological Sciences, 367, 2923–2934.

Real, L. A. (1977). The kinetics of functional response. The American Naturalist, 111, 289–300.

Rohr, T., Richardson, A. J., Lenton, A., Chamberlain, M. A. & Shadwick, E. H. (2023). Zooplankton grazing is the largest source of uncertainty for marine carbon cycling in CMIP6 models. Communications Earth & Environment, 4, 212.

Rohr, T., Richardson, A. J., Lenton, A. & Shadwick, E. (2022). Recommendations for the formulation of grazing in marine biogeochemical and ecosystem models. Progress in Oceanography, 208, 102878.

Rosenzweig, M. L. (1969). Why the prey curve has a hump. American Naturalist, 103, 81–87.

Rosenzweig, M. L. (1971). Paradox of enrichment: Destabilization of exploitation ecosystems in ecological time. Science, 171, 385–387.

Rosenzweig, M. L. & MacArthur, R. H. (1963). Graphical representation and stability conditions of predator-prey interactions. American Naturalist, 97, 209–223.

Roy, S. & Chattopadhyay, J. (2007). The stability of ecosystems: A brief overview of the paradox of enrichment. Journal of Biosciences, 32, 421–428.

Rubin, J. E., Earn, D. J. D., Greenwood, P. E., Parsons, T. L. & Abbott, K. C. (2023). Irregular population cycles driven by environmental stochasticity and saddle crawlbys. Oikos, 2023, e09290.

Seo, G. & Kot, M. (2008). A comparison of two predator–prey models with Holling’s type I functional response. Mathematical biosciences, 212, 161–179.

Seo, G. & Wolkowicz, G. S. K. (2015). Existence of multiple limit cycles in a predator-prey model with arctan(ax) as functional response. Communications in Mathematical Analysis, 18, 64–68.

Seo, G. & Wolkowicz, G. S. K. (2018). Sensitivity of the dynamics of the general Rosenzweig–MacArthur model to the mathematical form of the functional response: a bifurcation theory approach. Journal of Mathematical Biology, 76, 1873–1906.

Sjöberg, S. (1980). Zooplankton feeding and queueing theory. Ecological Modelling, 10, 215–225.

Skalski, G. T. & Gilliam, J. F. (2001). Functional responses with predator interference: Viable alternatives to the Holling Type II model. Ecology, 82, 3083–3092.

Stouffer, D. B. & Novak, M. (2021). Hidden layers of density dependence in consumer feeding rates. Ecology Letters, 24, 520–532.

Uiterwaal, S. F., Dell, A. I. & DeLong, J. P. (2018). Arena size modulates functional responses via behavioral mechanisms. Behavioral Ecology.

Uiterwaal, S. F. & DeLong, J. P. (2024). Foraging rates from metabarcoding: Predators have reduced functional responses in wild, diverse prey communities. Ecology Letters, 27, e14394.

Uiterwaal, S. F., Lagerstrom, I. T., Lyon, S. R. & DeLong, J. P. (2022). FoRAGE database: A compilation of functional responses for consumers and parasitoids. Ecology, 103, e3706.

Uszko, W., Diehl, S., Pitsch, N., Lengfellner, K. & Müller, T. (2015). When is a type III functional response stabilizing? Theory and practice of predicting plankton dynamics under enrichment. Ecology, 96, 3243–3256.

Volterra, V. (1926). Fluctuations in the abundance of a species considered mathematically. Nature, 118, 558–560.

Wootton, J. T. & Emmerson, M. (2005). Measurement of interaction strength in nature. Annual Review of Ecology, Evolution, and Systematics, 36, 419–444.

