## Supplementary material for "In defense of the original Type I functional response: The frequency and population-dynamic effects of feeding on multiple prey at a time": SOM

#### Table of Contents

**Multi-prey functional response model** **S1**
- Derivations . . . S1
- Proportion of predators feeding on 1 to $n$ prey . . . S4
- Equivalence of eqns. 4 and 5 for integer values of $n$ . . . S5

**Analysis of FoRAGE datasets** **S6**
- Data exclusions and re-scaling . . . S6
- Supplementary figures and statistical tables . . . S7

**Population-dynamic effects** **S12**
- Supplementary figures . . . S12

**A reformulation of the extended Steady State Saturation model** **S13**

### Multi-prey functional response model

#### Derivations

More explicit derivations of the Type II and multi-prey models are as follows.

##### Holling Type II model

Assuming a predator population $P$ of fixed size that is composed of only $P_S$ searching and $P_H$ handling sub-populations, let the rate of change in abundance of the two sub-populations be described by

$$\frac{dP_S}{dt} = -aNP_S + \tfrac{1}{h}P_H \tag{S.1a}$$

$$\frac{dP_H}{dt} = aNP_S - \tfrac{1}{h}P_H\,. \tag{S.1b}$$

Correspondingly, the rate at which eaten prey $N_e$ are generated is

$$\frac{dN_e}{dt} = \tfrac{1}{h}P_H\,. \tag{S.2}$$

As in the main text, $a$ is the per capita attack rate, $h$ the handling time, and $N$ the prey's abundance (which is also assumed fixed at the behavioral time scale we are considering).

Setting $\frac{dP_H}{dt} = 0$ (i.e. assuming steady state conditions), we substitute $(P - P_H)$ for $P_S$ and rearrange to determine the proportion of the whole population that is busy handling:

$$aN(P - P_H) = \tfrac{1}{h}P_H \tag{S.3a}$$

$$\implies aNP = aNP_H + \tfrac{1}{h}P_H \tag{S.3b}$$

$$= (aN + \tfrac{1}{h})P_H \tag{S.3c}$$

$$\implies \frac{P_H}{P} = \frac{aN}{aN + \frac{1}{h}} \tag{S.3d}$$

$$= \frac{ahN}{1 + ahN}\,. \tag{S.3e}$$

The total number of handling predators is thus

$$P_H = \frac{ahNP}{1 + ahN}\,. \tag{S.4}$$

Since the rate at which each of these $P_H$ predators finishes handling its prey is $\frac{1}{h}$, it follows that the rate at which eaten prey are "generated" by the whole predator population is

$$\frac{dN_e}{dt} = \frac{1}{h}P_H = \frac{aNP}{1 + ahN} \tag{S.5}$$

and thus that the *per predator* feeding rate (the functional response) is

$$f(N) = \frac{1}{P}\frac{dN_e}{dt} = \frac{1}{h}\frac{P_H}{P} = \frac{aN}{1 + ahN}\,. \tag{S.6}$$

##### Multi-prey model

Again assume a predator population $P$ of fixed size that is composed of $P_S$ searching and handling sub-populations, but now split handling predators into those capable of searching while handling less than $n$ prey individuals at any moment time. We therefore have that

$$P = P_S + P_{H_1} + P_{H_2} + \ldots + P_{H_n} \tag{S.7}$$

and describe the rate of change for each sub-populations by

$$\begin{aligned}
\frac{dP_S}{dt} &= -aNP_S + \tfrac{1}{h}P_{H_1} && \text{(S.8a)}\\
\frac{dP_{H_1}}{dt} &= aNP_S - \tfrac{1}{h}P_{H_1} && \text{(S.8b)}\\
\frac{dP_{H_2}}{dt} &= aNP_{H_1} - \tfrac{1}{h}P_{H_2} && \text{(S.8c)}\\
&\vdots\\
\frac{dP_{H_n}}{dt} &= aNP_{H_{(n-1)}} - \tfrac{1}{h}P_{H_n}\,. && \text{(S.8d)}
\end{aligned}$$

Correspondingly, the rate at which eaten prey $N_e$ are generated is now

$$\frac{dN_e}{dt} = \tfrac{1}{h}\sum_{i=1}^{n} P_{H_i}\,. \tag{S.9}$$

By setting $\frac{dP_{H_i}}{dt} = 0$ for all sub-populations, rearranging, and iteratively substituting, we have that

$$\begin{aligned}
aNP_S = \tfrac{1}{h}P_{H_1} \implies P_{H_1} &= ahNP_S && \text{(S.10a)}\\
aNP_{H_1} = \tfrac{1}{h}P_{H_2} \implies P_{H_2} &= ahNP_{H_1} && \text{(S.10b)}\\
&= ahN(ahNP_S) && \text{(S.10c)}\\
&= (ahN)^2 P_S && \text{(S.10d)}\\
aNP_{H_2} = \tfrac{1}{h}P_{H_3} \implies P_{H_3} &= ahNP_{H_2} && \text{(S.10e)}\\
&= ahN((ahN)^2 P_S) && \text{(S.10f)}\\
&= (ahN)^3 P_S && \text{(S.10g)}\\
&\vdots\\
aNP_{H_{(n-1)}} = \tfrac{1}{h}P_{H_n} \implies P_{H_n} &= ahNP_{H_{(n-1)}} && \text{(S.10h)}\\
&= ahN((ahN)^{n-1} P_S) && \text{(S.10i)}\\
&= (ahN)^n P_S\,, && \text{(S.10j)}
\end{aligned}$$

with the last lines for $P_{H_n}$ inferred by induction. The proportional abundance of each $i$th sub-population is thus

$$\frac{P_{H_i}}{P} = \frac{(ahN)^i P_S}{P} \tag{S.11a}$$
$$= \frac{(ahN)^i P_S}{P_S + P_{H_1} + P_{H_2} + \ldots + P_{H_n}} \tag{S.11b}$$
$$= \frac{(ahN)^i P_S}{P_S + ahNP_S + \ldots + (ahN)^n P_S} \tag{S.11c}$$
$$= \frac{(ahN)^i}{1 + ahN + \ldots + (ahN)^n} \tag{S.11d}$$
$$= \frac{(ahN)^i}{1 + \sum_{k=1}^{n} (ahN)^k} \,. \tag{S.11e}$$

Each of the sub-populations generates eaten prey at rate $\frac{1}{h}$, thus the rate at which eaten prey are generated by the whole population is

$$\frac{dN_e}{dt} = \frac{1}{h} \sum_{i=1}^{n} P_{H_i} \tag{S.12a}$$
$$= \frac{1}{h} \sum_{i=1}^{n} \frac{P_{H_i}}{P} P \tag{S.12b}$$
$$= \frac{1}{h} \sum_{i=1}^{n} \frac{(ahN)^i}{1 + \sum_{k=1}^{n} (ahN)^k} P \tag{S.12c}$$
$$= \frac{\frac{1}{h} \sum_{i=1}^{n} (ahN)^i}{1 + \sum_{i=1}^{n} (ahN)^i} P \,. \tag{S.12d}$$

The *per predator* feeding rate is therefore

$$f(N) = \frac{1}{P} \frac{dN_e}{dt} = \frac{\frac{1}{h} \sum_{i=1}^{n} (ahN)^i}{1 + \sum_{i=1}^{n} (ahN)^i} \tag{S.13}$$

as given in eqn. 4 of the main text.

**Proportion of predators feeding on 1 to $n$ prey**

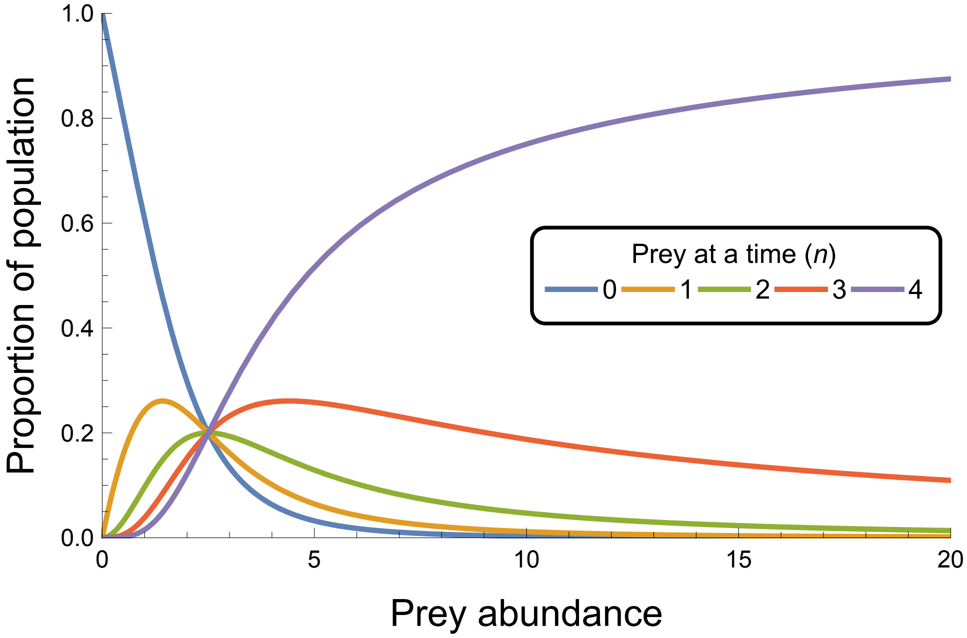


Figure S.1: The expected proportions of predator individuals that will be observed not feeding or handling $i = 1, 2, 3$ or $4$ prey changes with prey abundance (here visualized for a predator population whose individuals can handle up to $n = 4$ prey at a time). Individuals from each of the handling groups consumes prey at rate $1/h$, thus the predator population's (i.e. the average individual's) functional response is the product of $1/h$ and the sum of these handling-predator proportions. The prey abundance at which the expected proportions of individuals handling 0, 1, 2, 3 or 4 prey are all equal occurs at prey abundance $1/ah$. *Parameter values*: the attack rate is $a = 0.1$ and the handling time is $h = 4$.

##### Equivalence of eqns. 4 and 5 for integer values of $n$

Letting $n = 1$, we have

$$f(N) = \frac{aN(1-(ahN)^n)}{1-(ahN)^{n+1}} = \frac{aN(1-(ahN))}{1-(ahN)^2} = \frac{aN(1-ahN)}{1^2-(ahN)^2} = \frac{aN(1-ahN)}{(1+ahN)(1-ahN)} = \frac{aN}{1+ahN}\,.$$

Letting $n = 2$, we have

$$f(N) = \frac{aN(1-(ahN)^n)}{1-(ahN)^{n+1}} = \frac{aN(1-(ahN)^2)}{1-(ahN)^3} = \frac{aN(1+ahN)(1-ahN)}{(1+ahN+(ahN)^2)(1-ahN)} = \frac{aN(1+ahN)}{1+ahN+(ahN)^2} = \frac{\frac{1}{h}\sum_{i=1}^{2}(ahN)^i}{1+\sum_{i=1}^{2}(ahN)^i}\,.$$

Letting $n = 3$, we have

$$f(N) = \frac{aN(1-(ahN)^n)}{1-(ahN)^{n+1}} = \frac{aN(1-(ahN)^3)}{1-(ahN)^4} = \frac{aN(1+ahN+(ahN)^2)(1-ahN)}{(1+ahN+(ahN)^2+(ahN)^3)(1-ahN)} = \frac{aN(1+ahN+(ahN)^2)}{1+ahN+(ahN)^2+(ahN)^3)} = \frac{\frac{1}{h}\sum_{i=1}^{3}(ahN)^i}{1+\sum_{i=1}^{3}(ahN)^i}\,.$$

Their equivalence for higher integer values of $n$ follows similarly.

### Analysis of FoRAGE datasets

#### Data exclusions and re-scaling

The most recent version of FoRAGE (v.4) contains a total of 3013 datasets from which we excluded 422 for our analyses. Most of these were excluded because they entailed less than 4 prey-abundance treatment levels or because they had fewer than 15 data points (i.e. replicates) overall, but we also excluded several datasets because they provided prey abundances as densities for treatments that were implemented using arenas of varying size without specifying what those arena sizes were; entailed feeding rates of a variable but unspecified number of predators known to exhibit predator-dependent feeding rates; and/or entailed feeding rates of variable but unspecified experimental duration. Nine datasets were excluded because our models failed to reach convergence.

Our analyses required integer counts of prey abundance and eaten prey because we assumed binomial and Poisson likelihood functions to accommodate the increasing variance that accompanies an increase in the expected number of eaten prey (Novak & Stouffer, 2021b). For most datasets in which prey abundances were expressed as prey densities and/or predation was expressed as feeding rates, integer counts of prey abundance and prey eaten could be calculated using provided information on the area size(s) used (area or volume), the number of predators per treatment, and experimental duration(s). For raw-data datasets where this information was not provided, as well as datasets expressing densities and feeding rates on a mass basis (e.g., micro-grams of prey available or eaten), we (i) multiplied prey densities by the minimum scalar value necessary to obtain integer values across all prey densities (which we then used as prey abundance counts), and (ii) multiplied prey feeding rates by the minimum scalar value necessary to obtain integer values across all feeding rates (which we then used as counts of prey eaten). We multiplied prey abundances by an additional minimum scalar value for non-replacement datasets (reported as raw-data or as means) where the units in which densities and feeding rates were measured caused there to be more prey eaten than were seemingly available. Although these procedures will have altered the interpretation of the attack rate and handling time parameters (i.e. our estimates of $a$ and $h$ are not comparable across datasets), neither procedure will have affected our estimates of $n$ for the multi-prey model (because it is dimensionless) except, potentially, through an influence on the variance of the likelihood models (larger counts of prey eaten being permitted a higher variance than low counts of prey eaten). Although we did not observe any relationship between estimates of $n$ and the magnitudes of re-scaling across our datasets, its potential influence is worthy of future analytical study.

#### Penalized likelihood

Many datasets were not sufficiently informative to constrain estimates of $n$ and $\phi$. We therefore implemented a penalized likelihood approach, augmenting the two aforementioned likelihood functions with a penalty term proportional to the values of $n$ and $\phi$ to discourage large values of $n$ and $\phi$. More specifically, we performed model fitting using

$$\mathcal{L}_p = \mathcal{L} + \lambda \cdot \ln(n) + \lambda \cdot \ln(\phi) \tag{S.14}$$

as the loss function, where $\mathcal{L}$ is the negative log-likelihood and $\lambda$ determines the strength of the penalty for values of $n$ and $\phi$. Although it is possible to treat $\lambda$ as a free parameter that is estimated for each dataset, we set $\lambda = 1/\ln(20)$. A value of $n$ or $\phi$ equal to 20 therefore penalized the negative log-likelihood by 1 unit (equivalent to 1/2 the penalty associated with each parameter of a model under AIC).

#### Supplementary figures and statistical tables

(a) Insects

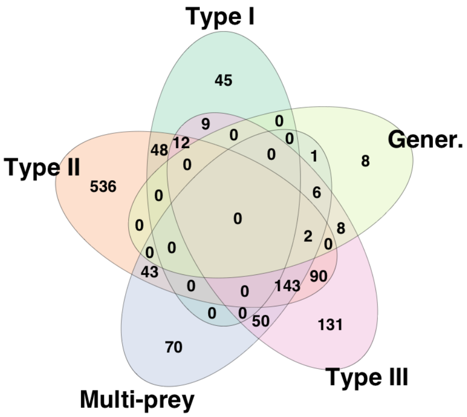


(b) Arachnids

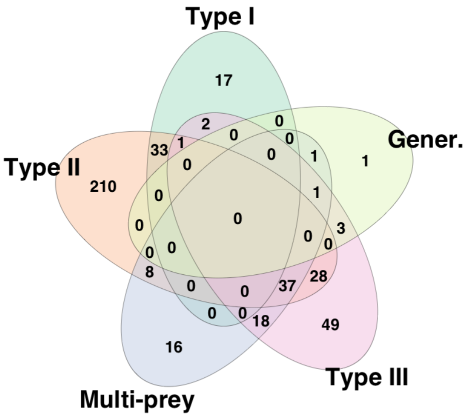


(c) Crustaceans

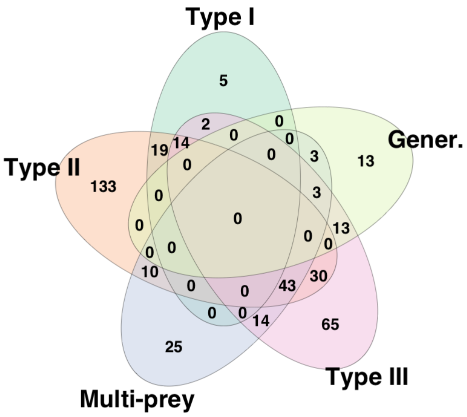


(d) Fishes

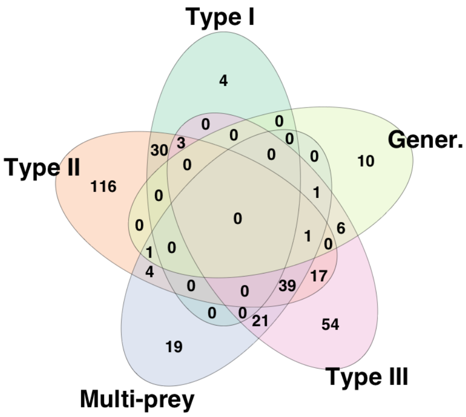


Figure S.2: Venn diagrams categorizing the datasets of the four most common predator groups by their support for one or more of the considered models based on their BIC scores.

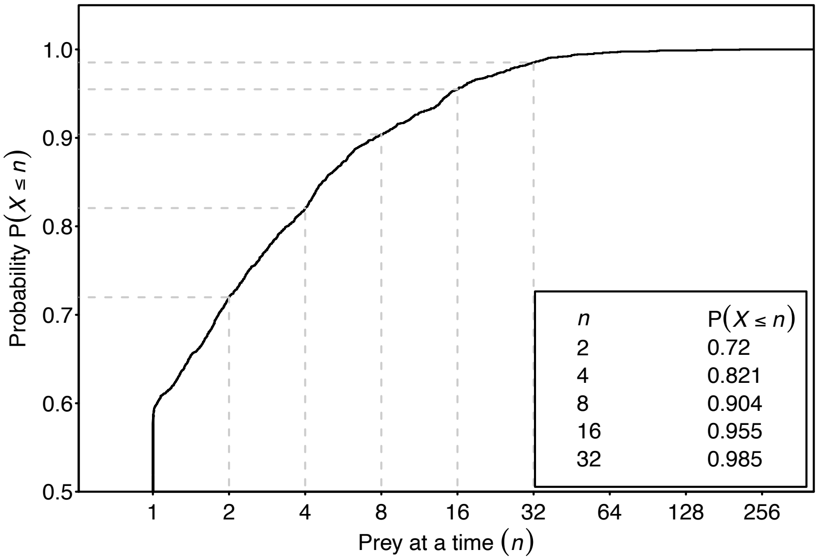


Figure S.3: Cumulative probability distribution of the estimates of $n$ (assuming the multi-prey model) from across all datasets excluding those for which the Type I model alone performed best.

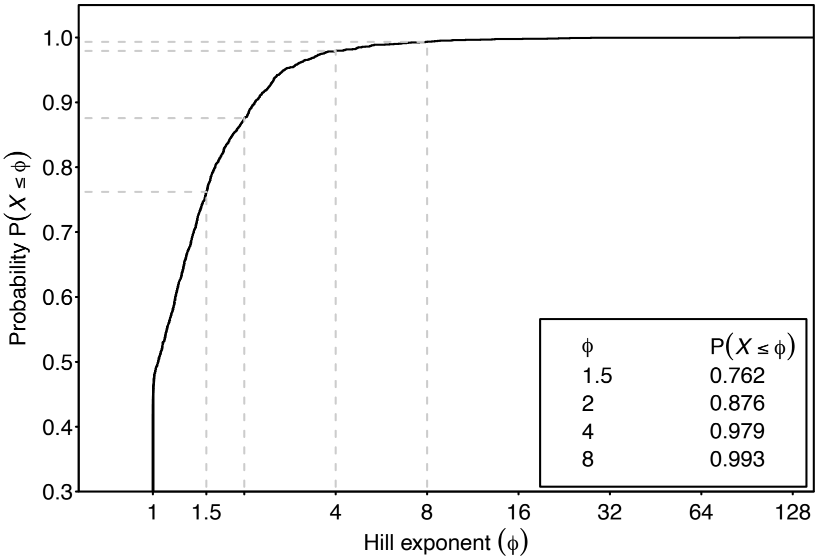


Figure S.4: Cumulative probability distribution of the estimates of $\phi$ (assuming the Holling-Real Type III model) from across all datasets excluding those for which the Type I model alone performed best.

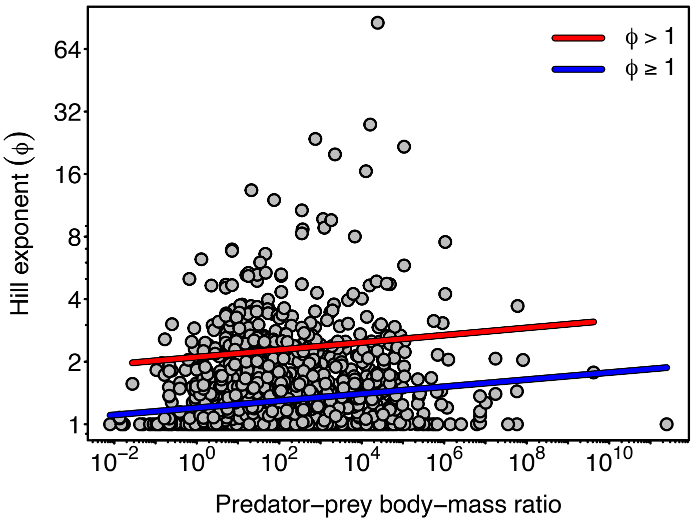


Figure S.5: The relationship between $log_2(\phi)$ and $log_{10}$(PPMR) assuming the Holling-Real model excluding datasets for which the Type I model alone performed best (Table S.2).

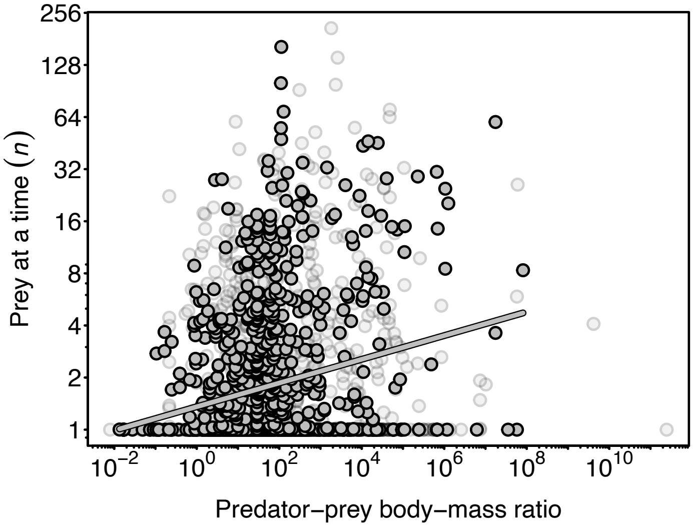


Figure S.6: The relationship between $log_2(n)$ and $log_{10}$(PPMR) assuming the multi-prey model when considering only those datasets having a sample size greater than the median sample size of all datasets excluding those for which the Type I model alone performed best (Table S.3).

Table S.1: Summary statistics (with 95% confidence intervals) for the least-squares linear regressions of $log_2(n)$ of the multi-prey model on $log_{10}$(PPMR) when considering all studies ($n \geq 1$) or only those studies for which $n$ >1.

| | Estimates | |
|---|---|---|
| | $n \geq 1$ | $n$ >1 |
| Intercept | 0.546*** (0.455, 0.638) | 1.976*** (1.806, 2.147) |
| $log_{10}$(PPMR) | 0.145*** (0.106, 0.184) | 0.190*** (0.122, 0.258) |
| Observations | 2,137 | 715 |
| $R^2$ | 0.024 | 0.041 |
| Adjusted $R^2$ | 0.024 | 0.039 |
| Residual Std. Error | 1.342 (df = 2135) | 1.334 (df = 713) |
| F Statistic | 53.006*** (df = 1; 2135) | 30.186*** (df = 1; 713) |

*p<0.1; **p<0.05; ***p<0.01

Table S.2: Summary statistics (with 95% confidence intervals) for the least-squares linear regressions of $log_2(\phi)$ of the Holling-Real Type III on $log_{10}$(PPMR) when considering all studies ($\phi \geq 1$) or only those studies for which $\phi$ >1.

| | Estimates | |
|---|---|---|
| | $\phi \geq 1$ | $\phi$ >1 |
| Intercept | 0.262*** (0.222, 0.302) | 1.074*** (0.974, 1.173) |
| $log_{10}$(PPMR) | 0.056*** (0.039, 0.073) | 0.058*** (0.020, 0.097) |
| Observations | 2,137 | 511 |
| $R^2$ | 0.020 | 0.017 |
| Adjusted $R^2$ | 0.019 | 0.015 |
| Residual Std. Error | 0.583 (df = 2135) | 0.667 (df = 509) |
| F Statistic | 42.597*** (df = 1; 2135) | 8.810*** (df = 1; 509) |

*p<0.1; **p<0.05; ***p<0.01

Table S.3: Summary statistics (with 95% confidence intervals) for the least-squares linear regression of $log_2(n)$ of the multi-prey model on $log_{10}$(PPMR) when considering only those studies having a sample size greater than the median sample size of all studies.

| | Sample size >36 |
|---|---|
| Intercept | 0.440*** (0.309, 0.571) |
| $log_{10}$(PPMR) | 0.228*** (0.167, 0.289) |
| Observations | 981 |
| $R^2$ | 0.052 |
| Adjusted $R^2$ | 0.051 |
| Residual Std. Error | 1.289 (df = 979) |
| F Statistic | 53.442*** (df = 1; 979) |

*p<0.1; **p<0.05; ***p<0.01

Table S.4: Summary statistics (with 95% confidence intervals) for the multiple least-squares linear regression of $log_2(n)$ of the multi-prey model on $log_{10}$(PPMR) × predator group for the four most common predator taxonomic groups.

| | Focal predators |
|---|---|
| Intercept (Insect) | 0.544*** (0.409, 0.678) |
| $log_{10}$(PPMR) | 0.167*** (0.090, 0.244) |
| Arachnid | −0.305** (−0.580, −0.029) |
| Crustacean | 0.208 (−0.063, 0.479) |
| Fish | −0.164 (−0.680, 0.352) |
| $log_{10}$(PPMR):Arachnid | 0.269** (0.050, 0.488) |
| $log_{10}$(PPMR):Crustacean | −0.061 (−0.165, 0.044) |
| $log_{10}$(PPMR):Fish | −0.029 (−0.201, 0.144) |
| Observations | 1,917 |
| $R^2$ | 0.032 |
| Adjusted $R^2$ | 0.029 |
| Residual Std. Error | 1.318 (df = 1909) |
| F Statistic | 9.150*** (df = 7; 1909) |

*p<0.1; **p<0.05; ***p<0.01

#### Population-dynamic effects

##### Supplementary figures

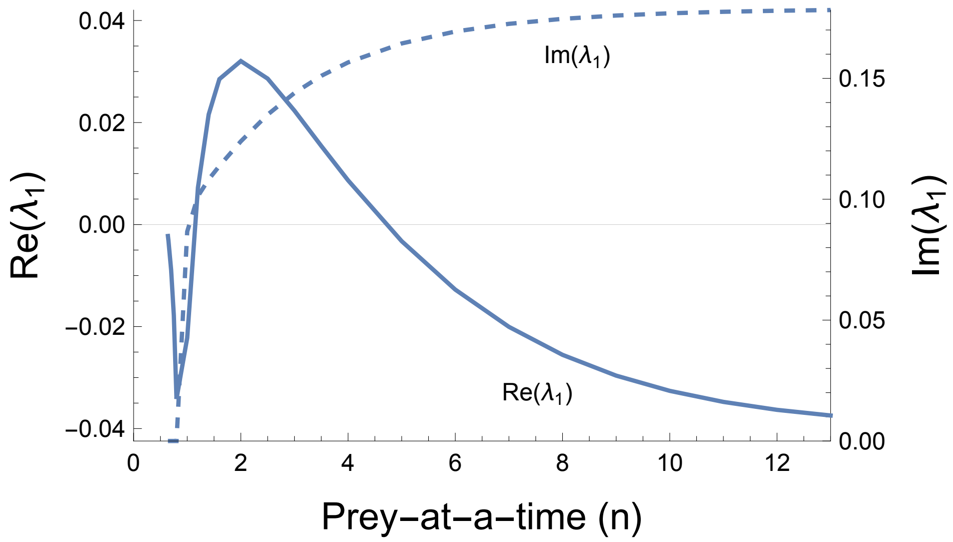


Figure S.7: The coexistence state is asymptotically stable when the real part of the dominant eigenvalue Re($\lambda_1$) is negative. This occurs for $n \approx 1$ where it is globally stable and for $n > 5$ where it is only locally stable. Post-perturbation dynamics towards the stable equilibrium exhibit monotonic damping when the imaginary part Im($\lambda_1$) is zero as occurs for $n \approx 1$, but exhibit damped oscillations when Im($\lambda > 0$) as occurs for higher $n$. *Other parameter values* as in Fig. 3.

#### A reformulation of the extended Steady State Saturation model

Jeschke *et al.* (2004) introduced a functional response model that, like the multi-prey model, is capable of exhibiting a continuum of shapes between the Type I and Type II response forms. In its original formulation, their model is written as

$$\frac{e(1 + aN(b + c)) - \sqrt{e\left(4acN + e(1 + aN(b - c))^2\right)}}{2c(e(1 + abN) - 1)}, \tag{S.15}$$

where $N$ is the prey's abundance, $a$ is the attack rate, $b$ is the handling time, $c$ is the digestion time, and $e$ is a dimensionless shape parameter interpreted as affecting the trade-off between search effort and hunger level (i.e. gut fullness). The model approaches the rectilinear model as $e \to \infty$ when $b = 0$ (see Fig. A2 of Jeschke *et al.*, 2004). For $e = 1$ it reduces to the "Steady State Saturation" (SSS) model of Jeschke *et al.* (2002), written in its original formulation as

$$\frac{1 + aN(b + c) - \sqrt{1 + aN\left(2(b + c) + aN(b - c)^2\right)}}{2abcN}. \tag{S.16}$$

Both models may be expressed in a formulation more similar to the Holling form that eases a comparison to other functional response models. This may be done by deriving them using the citardauq formula. The SSS may thereby be rewritten as

$$\frac{2aN}{1 + aN(b + c) + \sqrt{1 + aN\left(2(b + c) + aN(b - c)^2\right)}}. \tag{S.17}$$

(Note that the equation presented in the original version of Novak & Stouffer (2021a) is incorrect but has subsequently been corrected (Novak & Stouffer, 2024).) The extended SSS with parameter $e$ may be rewritten as

$$\frac{2aN}{1 + aN(b + c) + \frac{1}{e}\sqrt{e\left(4acN + e(1 + aN(b - c))^2\right)}}. \tag{S.18}$$

With four parameters, the extended SSS model is capable of exhibiting more variation in shape than the three-parameter multi-prey model. In particular, with sufficiently high $e$ and appropriately chosen non-zero values of $b$ and $c$, it exhibits curvature at the low prey abundances where the multi-prey model with high $n$ is effectively linear (see Figs. A1 and A2 of Jeschke *et al.*, 2004).
